# CATaN maps gene regulatory programs that shape genetic risk across complex diseases

**DOI:** 10.64898/2026.06.28.735120

**Authors:** Haruka Takahashi, Hiroaki Hatano, Michihiro Kono, Koma Haruta, Masahiro Nakano, Reza Bagherzadeh, Maximilian M. Drees, Yuki Oguma, Daichi Harita, Tsugumi Kawashima, Takahiro Arakawa, Hajime Inokuchi, Takahiro Nishino, Kenichiro Asahara, Takahiro Itamiya, Jun Inamo, Bunki Natsumoto, Yumi Tsuchida, Shuji Sumitomo, Akari Suzuki, Yuta Kochi, Keishi Fujio, Kazuhiko Yamamoto, Tazro Ohta, Eiryo Kawakami, Kazuyoshi Ishigaki

## Abstract

Causal variants of complex traits are enriched at transcription factor (TF) binding sites and are thought to contribute to pathology by disrupting TF activity and thereby causing transcriptome dysregulation. However, existing approaches typically address TF-mediated gene regulatory networks (TF-GRNs) and transcriptomes separately, and methods that jointly leverage both to systematically assess disease heritability remain limited. We aimed to develop a framework that jointly leverages TF-GRNs and transcriptomes to assess disease heritability. Here, we constructed a matrix encoding TF-GRNs and developed an unsupervised analytical pipeline, Canonical correlation Analysis of Transcriptome and TF-gene regulatory Networks (CATaN). CATaN applies canonical correlation analysis (CCA) to extract canonical correlation (CC) components, i.e., shared variation components between transcriptomes and TF-GRNs, and converts them into genome-wide functional annotation scores connected to stratified LD score regression (S-LDSC) for heritability analysis. We applied CATaN to eight datasets, including 19,198 bulk samples and 611,772 single cells from human and mouse sources, identifying 588 CC components that are significantly enriched for SNP heritability across 69 complex traits. Notably, functional annotation tracks based on these TF-GRNs are distinct from transcriptome signatures prioritized by LDSC-SEG, with greater heritability enrichment for a subset of traits. Finally, we suggest that CATaN may help prioritize candidate causal variants for experimental fine-mapping using genome editing. Together, integrating TF-GRNs with transcriptomes reveals disease-relevant regulatory programs that are not fully captured by transcriptome-based analyses alone.

## Introduction

Genome-wide association studies (GWASs) indicate that complex diseases arise from the cumulative effects of numerous causal variants^1,2,3^. Methods for partitioning SNP heritability, such as stratified LD score regression (S-LDSC)^4^, quantify the enrichment of SNP heritability within defined genomic functional regions. Analyses of publicly available TF chromatin immunoprecipitation sequencing (ChIP-seq) datasets^5^ have revealed that disease-associated heritability is concentrated within TF binding sites, for example, binding sites of NF-κB family members in rheumatoid arthritis (RA)^6^. These observations suggest that genetic variants perturb TF-mediated regulatory programs and that TF gene regulatory networks (TF-GRNs; defined here as the set of regulatory relationships between TFs and their putative target genes) represent an important mechanistic layer linking genetic variation to disease. Meanwhile, the growing availability of large-scale transcriptomic resources, including bulk and single-cell RNA sequencing datasets spanning diverse tissues and biological conditions, provides unprecedented opportunities to identify cellular programs relevant to disease genetics.

Several approaches have been developed to integrate either TF-GRNs or transcriptomic variation with GWAS signals. On the TF-GRN side, S-LDSC has been applied to TF ChIP-seq–derived annotations to partition heritability across regulatory elements^6^. IMPACT^7^ integrates TF motif information with epigenomic features, including ChIP-seq and chromatin accessibility data, to define cell-state-specific regulatory annotations for heritability enrichment analysis. More recently, Perturb-multiome has enabled experimental reconstruction of TF-GRNs through combinatorial CRISPR perturbation screens coupled with joint single-cell transcriptomic and chromatin accessibility profiling^8^. However, these approaches primarily focus on defining regulatory elements or characterizing regulatory network architecture rather than identifying transcriptomic programs enriched for disease heritability.

On the transcriptome side, the most widely used framework is LD score regression applied to specifically expressed genes (LDSC-SEG)^9^, which tests for enrichment of causal variants near genes exhibiting label-specific expression patterns. sc-linker^10^ extends this framework to single-cell RNA sequencing (scRNA-seq) data by deriving gene programs and linking them to SNP annotations through tissue-specific enhancer–gene maps before applying S-LDSC. Although these approaches have successfully highlighted disease-relevant cell types and cellular states, they rely primarily on transcriptomic variation and do not explicitly model TF-centered regulatory programs. Consequently, they cannot directly assess whether disease heritability is concentrated within transcriptional programs coordinated by specific TF regulatory networks.

A second limitation of existing transcriptome-based approaches is their reliance on predefined cellular annotations. Methods such as LDSC-SEG and sc-linker typically require investigators to define cell types, cellular states, or comparison groups a priori, potentially overlooking regulatory programs that transcend conventional cell-type boundaries. Several efforts have sought to reduce reliance on predefined annotations. One such approach is scDRS^11^, which scores individual cells based on the aggregate expression of GWAS-prioritized gene sets derived using MAGMA^12^. While powerful for identifying disease-relevant cells, scDRS aggregates genetic associations at the gene level and therefore does not explicitly model SNP-level genetic architecture. Another recent approach leverages paired snRNA-seq and snATAC-seq profiling to define annotations based on chromatin accessibility patterns associated with transcriptomic variation, followed by heritability enrichment analysis via S-LDSC^13^. Although this strategy captures cell-state-dependent regulatory elements without relying on predefined labels, it requires multimodal datasets, limiting its applicability relative to the substantially larger collection of single-modality scRNA-seq datasets.

Together, these limitations highlight an unmet need for a framework that integrates TF-GRNs with transcriptomic variation, operates without predefined cellular annotations, and directly leverages genome-wide SNP-level genetic information. To address this challenge, we developed CATaN (**<u>C</u>**anonical correlation **<u>A</u>**nalysis of **<u>T</u>**ranscriptome **<u>a</u>**nd TF-gene regulatory **<u>N</u>**etworks), an unsupervised framework that identifies shared variation between transcriptomes and TF-GRNs through canonical correlation analysis (CCA)^14^, producing genome-wide, single-base-resolution scores that reflect the regulatory potential underlying this shared variation. Although CCA has previously been used to identify shared variance between upstream regulatory sequence motifs and gene expression profiles^15^, to our knowledge it has not been applied to integrate TF ChIP-seq-derived regulatory networks with transcriptomic data for heritability enrichment analysis. Moreover, unlike the sequence motifs used in that earlier work, the ChIP-seq-derived TF-GRNs employed by CATaN capture cell-type-specific TF occupancy, providing complementary regulatory information^16^. By integrating TF-GRN information with transcriptomic variation, CATaN identified disease-relevant regulatory programs and heritability enrichments that had not been captured by existing transcriptome-only approaches.

## Results

### Overview of CATaN

Briefly, CATaN consists of three steps (Figure 1A):

1. Construct a TF-GRN matrix (genes × TF ChIP–seq samples) using exponential weighted TF-gene connectivity scores, with a half-decay distance of 10 kb selected to maximize how strongly CCA captures shared variation between the transcriptome matrix and the TF-GRN matrix; this distance captures regulatory contributions extending beyond proximal promoters, including enhancer-driven regulation (Methods; Supplementary Note; Figures S1B and S1C).
2. Construct a transcriptome matrix (genes × samples) with gene-expression values.
3. Apply CCA to link the two matrices and transform the CCA loading into genome-wide, single-base-pair-resolution scores.

**Figure 1.**
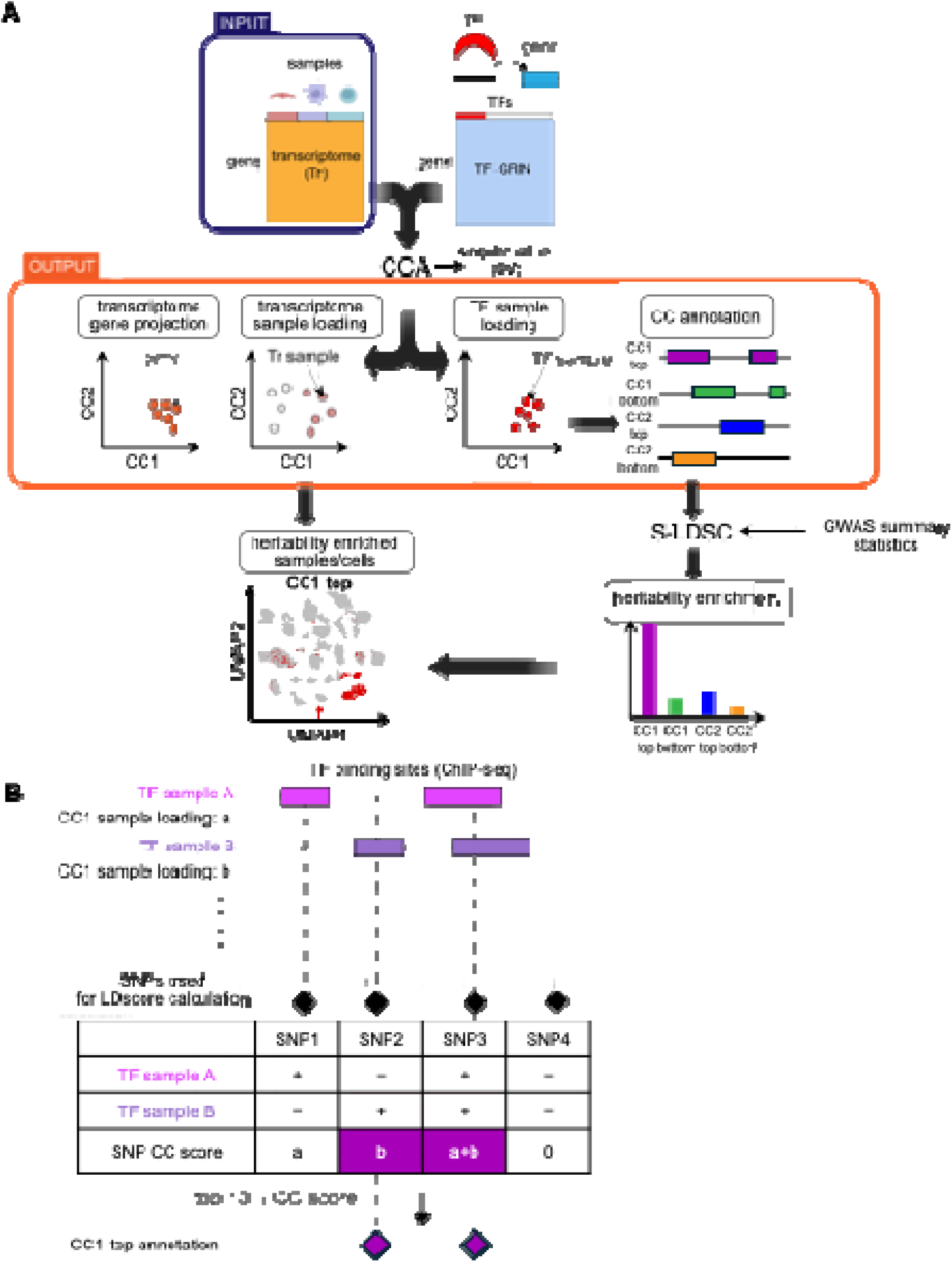
Overview of CATaN. (A) Schematic of Canonical correlation Analysis of Transcriptome and TF-gene regulatory Networks (CATaN). CATaN takes two input matrices—a transcriptome matrix (genes × samples) and a TF-Gene Regulatory network (TF-GRN) matrix (genes × TF ChIP-seq samples)—and applies canonical correlation analysis (CCA) to yield three outputs per canonical component (CC): transcriptome sample loadings, TF sample loadings and transcriptome gene projections. Singular values (SVs) serve as an indicator of CCA model fitness. TF sample loadings are converted into genome-wide CC annotation tracks using ChIP-seq binding profiles and combined with genome-wide association study (GWAS) summary statistics for SNP-heritability enrichment analysis. Heritability-enriched samples and cell are then identified by combining transcriptome sample loadings and heritability enrichment results. (B) Generation of CC annotation tracks. TF sample loadings are assigned to TF peak regions. For each SNP used in linkage disequilibrium (LD) score regression, scores from all TF peaks overlapping the SNP are summed, while SNPs outside any peak receive a score of 0. Binary annotations are then created by assigning a value of 1 to SNPs in the top or bottom 10% of the score distribution and 0 to all others. Created with BioRender.com.

We constructed a TF-GRN matrix by curating and processing a large TF ChIP-seq compendium comprising 2,868 profiles covering 410 TFs across 24 tissues and 287 cell types (Figure S1A and Table S1), and assembled transcriptome matrices spanning diverse biological contexts. CATaN applies CCA to identify shared variation between these two matrices. Applying CCA embeds transcriptome and TF ChIP-seq samples into a shared canonical space, yielding three outputs per canonical component: (i) TF sample loadings, (ii) transcriptome sample loadings, and (iii) gene scores in the transcriptome embedding (transcriptome gene projections) (Supplementary Note; Figure S1D). Intuitively, CATaN linearly projects transcriptomic variation onto TF-GRN components to yield genome-wide annotation tracks. TF sample loadings can be converted into genome-wide annotation tracks using ChIP-seq binding profiles (CC annotation; Figure 1B and Methods), constituting the fourth output of CATaN. Briefly, the CC scores of each SNP is calculated as the sum of TF sample loadings of all TF peaks overlapping the SNP for the given CC. The top 10% and bottom 10% of SNPs by score were each used to define a separate binary annotation. These CC annotations are used to test heritability enrichment and to prioritize candidate loci for downstream experimental characterization. Transcriptome sample loadings, together with heritability enrichment results, enable identification of the specific samples and cell types driving heritable disease risk. The singular values (SVs) of CCA serve as an indicator of model fit, reflecting how well each CC captures shared biological processes from the TF-GRN matrix and the transcriptome matrix.

### Directional consistency between TF and transcriptome matrices

CATaN identifies gene expression signatures that exhibit high co-variance across TF-GRN and transcriptome matrices, based on the assumption that genes directly regulated by a given transcription factor are predominantly up– or down-regulated in a coherent manner. We tested this assumption using three independent lines of evidence. We first evaluated a TF overexpression experiment. We overexpressed the NF-κB subunit RELA (also known as TF65) using a CRISPRa^17^ system and quantified gene expression by RNA-sequencing (RNA-seq) (see Figure 2A and Methods). Quality control analyses indicated that the CRISPRa perturbation was effective: CRISPRa increased RELA expression approximately 7.5-fold versus control (Figure 2B); Principal Component Analysis (PCA) showed clear separation of RELA-overexpressed and control samples along PC1 (Figure S2A); and principal variance components analysis (PVCA)^18^ estimated that sample condition accounted for ∼10% of total variance (Figure S2B). RELA overexpression induced approximately three times more up-regulated genes compared to down-regulated genes (47 up-regulated genes vs. 17 down-regulated genes), supporting our assumption (Figures 2C and D, Table S2A). Gene set enrichment analysis (GSEA)^19^ using Molecular Signatures Database (MSigDB)^20^ Hallmark gene sets revealed that multiple pathways were significantly enriched among upregulated genes in RELA-overexpressing cells, including the TNF-α signaling via NF-κB pathway, which served as a positive control for RELA-mediated transcriptional regulation. In contrast, down-regulated genes did not show significant pathway enrichment, consistent with RELA acting predominantly as a transcriptional activator in this context (Figure S2C, Table S2B).

**Fig. 2.**
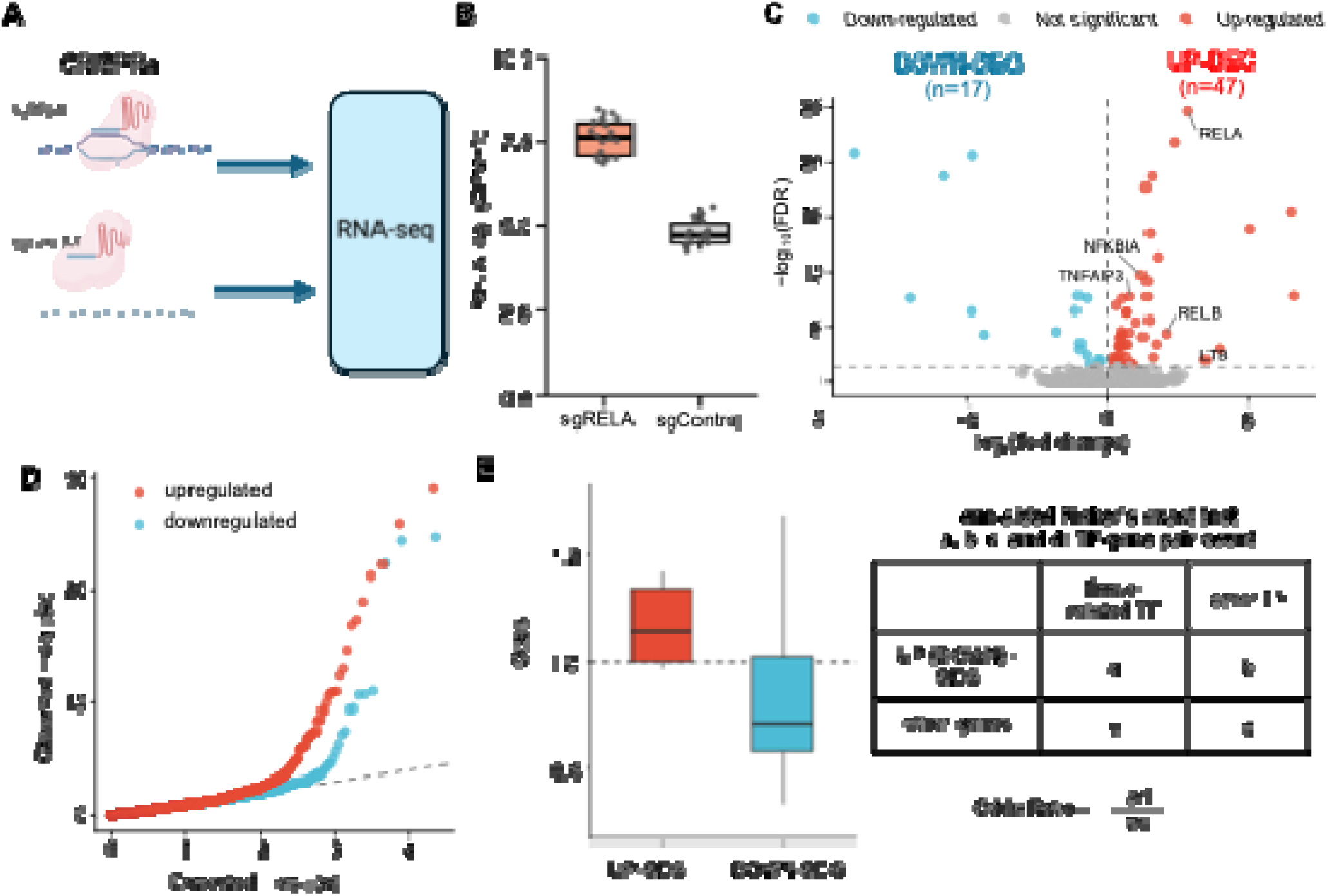
Concordant directional expression of TF-regulated genes. (A) Schematic of CRISPRa-mediated RELA activation and RNA-seq analysis. Jurkat cells stably expressing dCas9-VP64 were transduced with either a single guide RNA targeting RELA (sgRELA) or a non-targeting control (sgControl). (B) Box plots of RELA expression levels (log_2_ counts per million (CPM)) of sgRELA versus sgControl samples (*n* = 18 per group). (C) Volcano plot of differentially expressed genes (DEGs). Downregulated DEGs (*n* = 17) are shown in blue; upregulated DEGs (*n* = 47) in red. The dashed horizontal line indicates an FDR threshold of 0.05, and the vertical dashed line marks zero log_2_ (fold change). Gene names are labeled for significant DEGs involved in the NF-κB signaling pathway (KEGG: hsa04064). (D) Quantile–quantile (Q–Q) plot comparing the observed and expected −log₁₀(*p*) distributions of upregulated (red) and downregulated (blue) genes. The dashed line indicates *y = x*. (E) Box plots of odds ratios from one-sided Fisher’s exact tests assessing the enrichment of tissue-specific SEGs near tissue-associated TF binding sites versus other TF binding sites (*n* = 8). Odds ratios >1 indicate that SEGs are disproportionately found near tissue-related TFs. Upregulated and downregulated SEGs were tested separately.

We next evaluated tissue transcriptome data from GTEx^21^, in which tissue-defining TFs are well characterized. We assessed enrichment of tissue specifically expressed genes (SEGs) near tissue-associated TF binding sites (Figure 2E, Table S3A). Interestingly, tissue-specific TF binding sites were enriched in the upregulated tissue-SEGs compared to other TFs. This trend was less prominent in downregulated tissue-SEGs, with the exception of testis. This indicates that tissue-defining TFs bind preferentially near the genes they upregulate in each tissue, supporting the biological validity of the TF-GRN used in CATaN.

The same pattern was observed for clinical transcriptome data: binding sites of STAT and IRF family TFs, key mediators of the interferon response implicated in systemic lupus erythematosus (SLE)^22^, were enriched exclusively near up-regulated differentially expressed genes (DEGs) in SLE patients (mean log(odds) = 0.2), whereas, these binding sites were depleted near down-regulated DEGs (mean log(odds) = −0.2) (Figure S2D, Table S3B). Together, these results support a key assumption of CATaN, namely that TF-associated genes tend to show coherent directional changes in expression.

### Proof-of-concept analyses

We next evaluated CATaN from three complementary perspectives: (i) whether CATaN can correctly nominate the TFs driving transcriptome variation, (ii) whether CATaN is robust against overfitting, and (iii) whether CC annotation tracks recapitulate known gene–variant links. Using the RELA overexpression data described above (n = 6 each for sgRELA and sgControl), we constructed a transcriptome matrix comprising perturbed and non-perturbed samples and applied CATaN. Across 2,868 TF sample loadings, NF-κB ChIP-seq samples (n = 65) exhibited markedly higher CC1 loadings than other TFs (Figure 3A, Table S4; Fisher’s one-tailed test, odds ratio = 48; Bonferroni-corrected *p* = 1.68 × 10^−32^). Permutation of gene labels in the transcriptome data (Figure S3A) substantially reduced the Z-scored CC1 loadings of RELA (Figure 3B), indicating that the observed signal was not driven by technical inflation. These results indicate that CATaN captures shared variation between transcriptome and TF-GRN and can nominate the TFs driving transcriptome shifts.

**Fig. 3.**
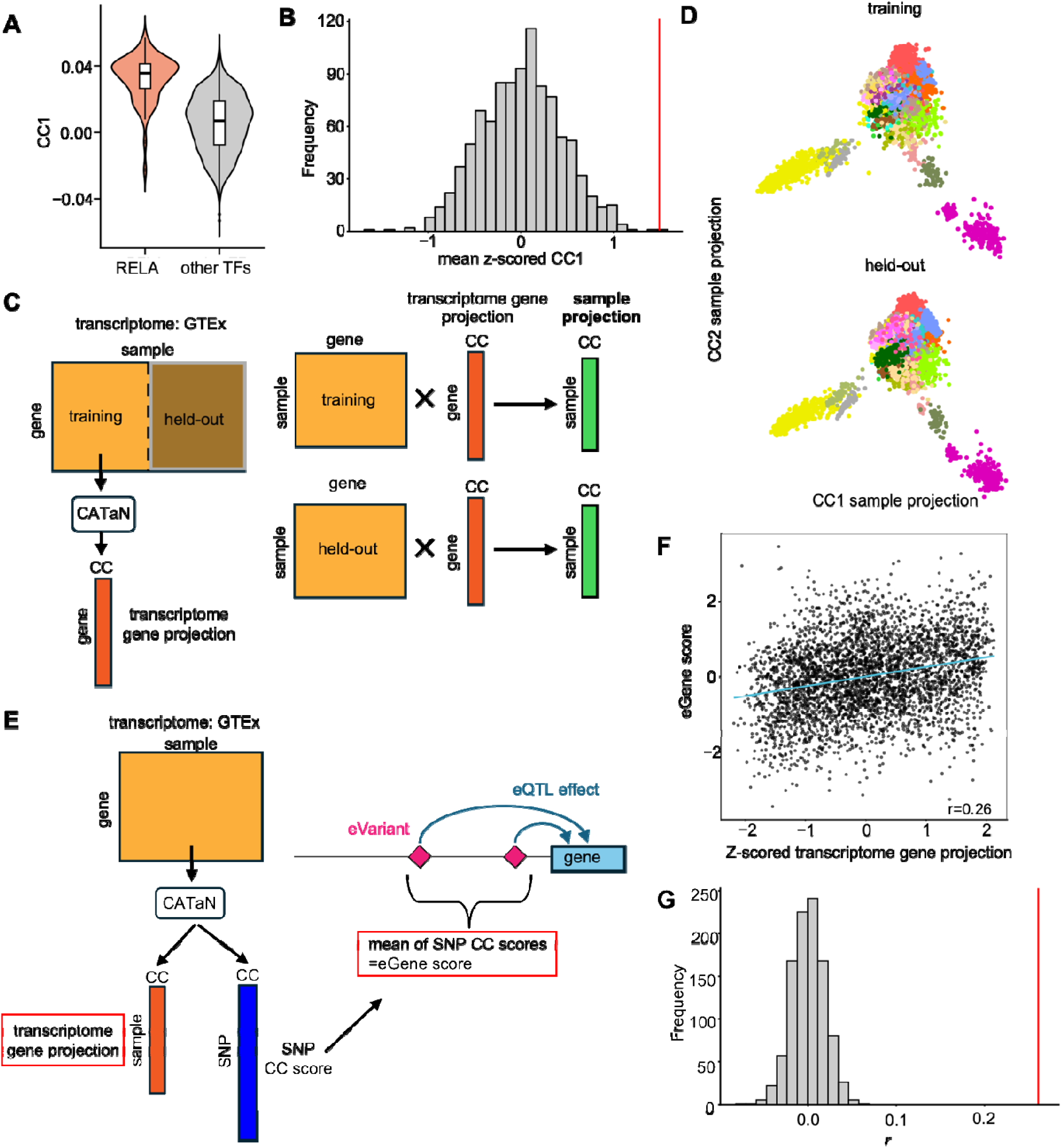
Proof-of-concept analyses. (A) Violin plots of CC1 TF sample loadings for NF-κB (RELA) ChIP-seq samples versus other TFs. CATaN was applied to CRISPRa RELA overexpression RNA-seq data (sgRELA and sgControl, *n* = 6 each). (B) Distribution of mean z-scored CC1 loadings for RELA from 1,000 permutations of the transcriptome matrix (CRISPRa RELA overexpression RNA-seq dataset). The red line indicates the observed value. (C) Schematic of holdout validation using GTEx transcriptome data. The GTEx transcriptome matrix was split into training (*n* = 4,277) and holdout sets (*n* = 4,278). CATaN was applied to the training set to derive transcriptome gene projections. The gene projections were then used as loadings to project both training and holdout samples onto the same CC space. (D) Scatter plots of training (top) and holdout (bottom) samples projected onto the CC1–CC2 space derived from training data, colored by tissue annotation. (E) Schematic of CC annotation track validation using fine-mapped eQTLs. For each eGene identified by fine-mapped *cis*-eQTLs from GTEx (v10), SNP-level CC scores of associated eVariants were averaged to define an eGene score. The correlation between transcriptome gene projections and eGene scores was then computed. (F) Scatter plot of transcriptome gene projections (x axis) versus inverse normal transformed (INT) eGene scores (y axis) for CC1. Each dot represents a gene (*n* = 3,646). *r*, Pearson correlation coefficient. The blue line indicates a linear regression fit. (G) Histogram of Pearson correlation coefficients (*r*) for CC1 between gene projections and eGene scores from null model in which gene labels in the transcriptome matrix were permuted before CCA (Figure S3A). The red line indicates the observed correlation.

To assess whether CATaN is robust against overfitting, we partitioned GTEx samples into two halves and independently applied CATaN to each subset. TF sample loadings from the two subsets were strongly correlated (Figure S3B). Moreover, when transcriptome gene projections derived from one subset (training subset) were used to project the held-out subset onto the same embedding space (Figure 3C), samples from the same tissue formed distinct clusters at corresponding positions (Figure 3D). Together, these results indicate that the latent space learned by CATaN reflects robust biological structure that generalizes across independent sample subsets, rather than overfitting.

Finally, we evaluated our CC annotation tracks against fine-mapped eQTLs from GTEx v10^23^, which provide independent gene–variant links. We applied CATaN to the GTEx transcriptome matrix and computed a per-gene CC score as the mean of variant-level CC scores of fine-mapped variants for each gene (the “eGene score”; Figure 3E). Gene projections positively correlated with eGene scores (*r* = 0.26 for CC1; Figure 3F), and this correlation was eliminated by permutation of the gene labels in the transcriptome matrix (Figure 3G), indicating that gene–variant connections inferred by CATaN are broadly concordant with eQTL mapping results.

### CATaN decomposes GTEx transcriptomic variation into biologically relevant axes

Having established that CATaN produces robust and interpretable outputs, we next examined whether its outputs provide coherent biologically relevant axes. GTEx is well suited for this purpose because its transcriptomic diversity and tissue-defining TFs are extensively characterized, enabling direct assessment of whether canonical axes capture known biology. CATaN embeds transcriptome and TF ChIP-seq samples into a shared canonical space, enabling joint characterization of gene expression and its upstream TF regulators. Each CC axis can be characterized through three CATaN outputs: transcriptome sample loadings (Tr-loading), TF sample loadings (TF-loading), and pathway enrichment of gene projections. We assessed the association between sample loadings and metadata using Fisher’s exact test with Bonferroni correction (see Methods). Tr-loadings revealed that brain tissues were enriched in CC1-top (odds ratio = 7.8 × 10^2^), whereas whole blood was enriched in CC1-bottom (odds ratio = 4.5 × 10^15^). CC2-bottom was enriched for whole blood (odds ratio = 2.0 × 10^2^) and spleen (odds ratio = 2.0 × 10^3^) (Figure 4A, Table S5A). TF-loading identified putative drivers for each axis: CC1-top was associated with REST (odds ratio = 75) and EZH2 (odds ratio = 7.1), both key regulators of neural gene programs; CC1-bottom with SPI1 (odds ratio = 18) and CC2-bottom with PAX5 (odds ratio = 32), a transcription factor essential for hematopoiesis and immune differentiation (Figure 4B, Table S5B). Pathway analyses based on gene projections further supported these patterns: genes included in the GABAergic synapse pathway were enriched in CC1-top (odds ratio = 9.3), whereas those in the NF-κB pathway were enriched in CC1-bottom (odds ratio = 6.5) and those in the B cell receptor pathway in CC2-bottom (odds ratio = 7.4) (Figure 4C, Table S5C). Together, these results demonstrate that CATaN decomposes transcriptomic heterogeneity into biologically relevant axes linking biological context (e.g., tissue, cell type, or disease status), TF activity, and gene programs.

**Fig. 4.**
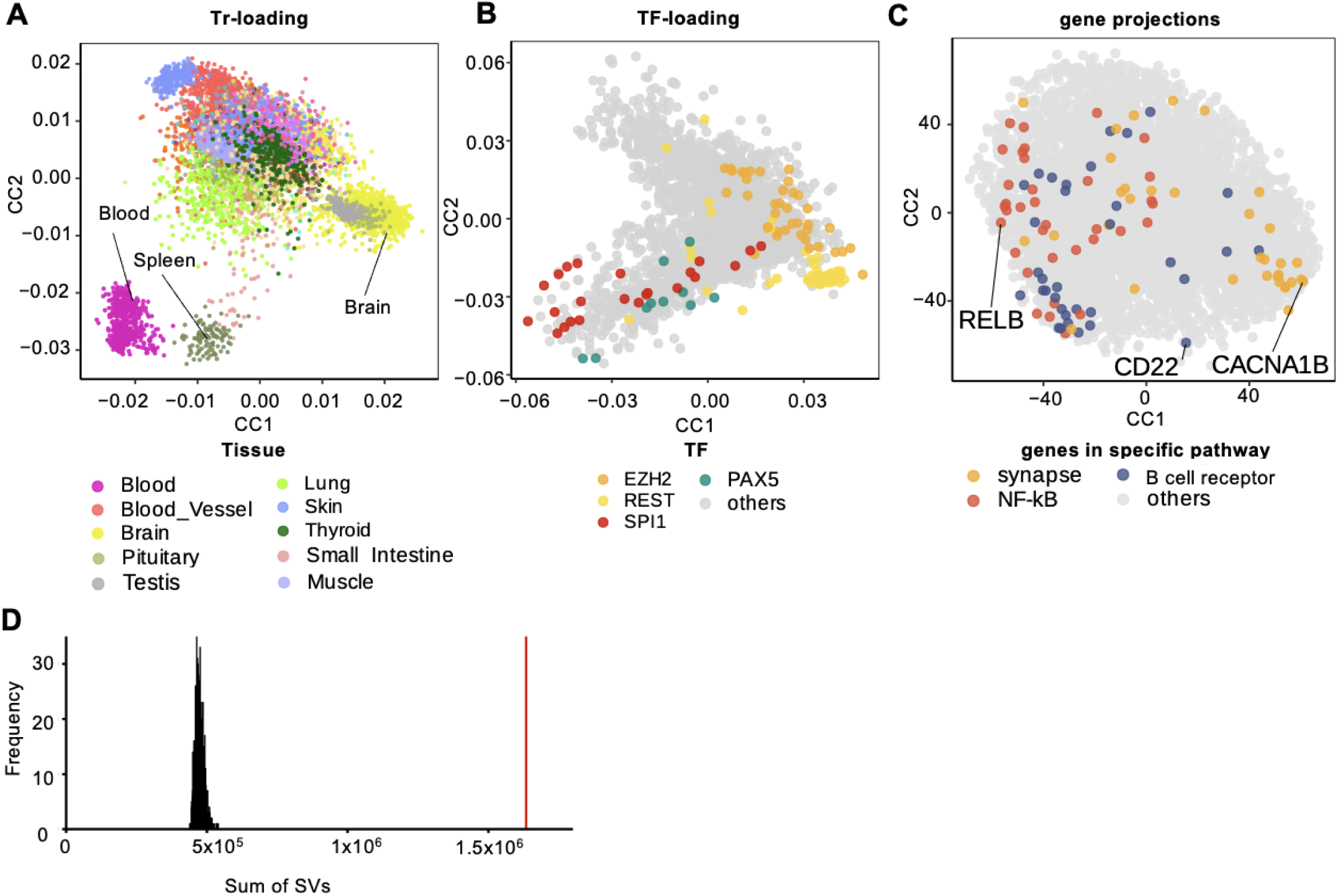
Evaluation of fitness and interpretation of CATaN output applied to GTEx. (A) Scatter plot of transcriptome sample loadings (Tr-loading) for CC1 and CC2, colored by GTEx tissu annotation (*n* = 8,555). Representative tissues are labeled and shown in the legend. (B) Scatter plot of TF sample loadings (TF-loading) for CC1 and CC2, colored by target TF in the TF-GRN matrix (*n* = 2,868). (C) Scatter plot of gene projections (*n* = 5,440) for CC1 and CC2, with genes belonging to selected KEGG pathways highlighted. ‘Synapse’ refers to the GABAergic synapse pathway. Genes with the highest absolute CC scores within each pathway are labeled. (D) Histogram of summed singular values (SVs) from CC1–CC10, computed from 1,000 permutations of the GTEx transcriptome matrix. The red line indicates the observed sum of singular values.

### CATaN application to multiple datasets

To assess the applicability of CATaN to multiple datasets, we applied it across eight transcriptome datasets spanning distinct biological contexts (Table 1): (i) bulk inter-tissue and cell-type dataset—GTEx (53 human autopsy tissues), human immune cell dataset (28 human immune cell types from healthy donors), and ImmGen^24,25^ (30 mouse immune cell types); (ii) single-cell datasets—one neural organoid and two immune-cell datasets; (iii) a stimulation dataset—bulk T cells under five conditions; and (iv) a clinical autoimmunity dataset—multi-autoimmune dataset (ten autoimmune diseases in the ImmuNexUT^26^ cohort).

**Table 1.** Datasets analyzed using CATaN. Summary of eight transcriptome datasets spanning distinct biological contexts, including data source, tissue or cell-type composition, modality, species and number of samples or cells. RNA-seq, RNA sequencing; scRNA-seq, single-cell RNA sequencing.

| name | source | Tissue/<br>cell-types | modality | species | The number of<br>samples/cells |
| --- | --- | --- | --- | --- | --- |
| GTEx | GTEx V6p | Multi tissues | RNA-seq | human | 8,555 |
| human immune<br>cell dataset | ImmuNexUT<br>Healthy control<br>(E-GEAD-397) | Multi cell-types | RNA-seq | human | 1,977 |
| ImmGen | ImmGen<br>(GSE15907) | Multi cell-types | microarray | mouse | 653 |
| ILC dataset | Skin-resident<br>innate lymphoid<br>cells<br>(GSE149622 ) | Multi cell-types | scRNA-seq | mouse | 26, 877 |
| RA synovium<br>dataset | AMP Phase 2<br>(syn52297840) | Multi cell-types | scRNA-seq | human | 576,600 |
| Neural<br>organoid<br>dataset | Cortical<br>organoids<br>(GSE163018) | Multi cell-types | scRNA-seq | human | 8,295 |
| Cytokine CD4<br>dataset | Original cytokine<br>stimulated CD4<br>RNA-seq | Single cell-type | RNA-seq | human | 138 |
| Multi-autoimmune<br>dataset | ImmuNexUT<br>10 diseases<br>(E-GEAD-397) | Single cell-type | RNA-seq | human | 9,852 |

We first evaluated the fitness of CATaN within each dataset from three perspectives: (i) whether SVs exceed those expected under a null model, (ii) whether TF sample loadings are associated with TF metadata, and (iii) whether transcriptome sample loadings are associated with sample metadata. SVs in the observed data were on average 3.5-fold higher than those from null data in GTEx (Figure 4D). Because CATaN is label-free and does not use TF sample metadata (410 TFs across 24 tissues), assessing how much of the variance in TF sample loadings is explained by this metadata provides an orthogonal validity check. Variance partitioning confirmed that TF identity (Figure S3C) and tissue each explained significantly non-zero variance in TF sample loadings (29% and 22%, respectively, in GTEx). Notably, compared with GTEx (across-tissue heterogeneity), human immune cell dataset and ImmGen (intra-immune cell heterogeneity) showed smaller tissue components and larger TF components (tissue 6.4% and TF 37% in human immune cell dataset, tissue 7.9% and TF 40% in ImmGen), indicating that CATaN metrics reflect the input data modality (Figure S3D, Table S6A). Reciprocally, transcriptome sample loadings also showed significant associations with metadata (Table S6B), further supporting the validity of CATaN. Together, these analyses show that CATaN recovers biologically meaningful covariation between TF-GRNs and diverse transcriptomes.

### Comparing CATaN-LDSC and LDSC-SEG

A key application of CATaN is estimating SNP-heritability enrichment via S-LDSC using CC annotation tracks. We first compared the genomic coverage of CC and SEG annotations. SEG (LDSC-SEG) annotations are transcriptome-based annotations in which a binary annotation is defined by extending a fixed 100-kb window around specifically expressed genes. In the GTEx dataset, SEG annotations encompassed 1.4–2.4 million SNPs (approximately 14–24% of all SNPs measured in the 1000 Genomes Project), whereas CC annotations were considerably narrower, covering approximately 1.0 million SNPs, as they were designed to capture 10% of all 1000 Genomes SNPs (Figure S4A).

To characterize the functional properties of these annotations, we compared the chromatin state composition of SNPs within each annotation using the Roadmap Epigenomics core 15-state ChromHMM model^27^ for E029 (primary monocytes from peripheral blood). Consistent with SEG annotations being defined around gene bodies, SNPs in SEG annotations were enriched for transcription-associated states (Wilcoxon rank-sum test; strong transcription: *p* = 1.3 × 10⁻□, weak transcription: *p* = 4.4 × 10⁻□) and weak repressed Polycomb states (*p* = 1.6 × 10⁻□), indicating that SEG annotations capture a substantial fraction of weakly repressed genomic regions. By contrast, since CC score is a linear combination of TF binding site annotations, CC annotations were enriched for enhancers (*p* = 3.3 × 10⁻□), active transcription start sites (*p* = 1.2 × 10⁻³), flanking active transcription start sites (*p* = 2.2 × 10⁻□), indicating that CC annotations preferentially capture regulatory elements involved in active transcription (Figure 5A). A similar trend was also observed using the 15-state ChromHMM model for E071 (brain) (Figure S4B).

**Fig. 5.**
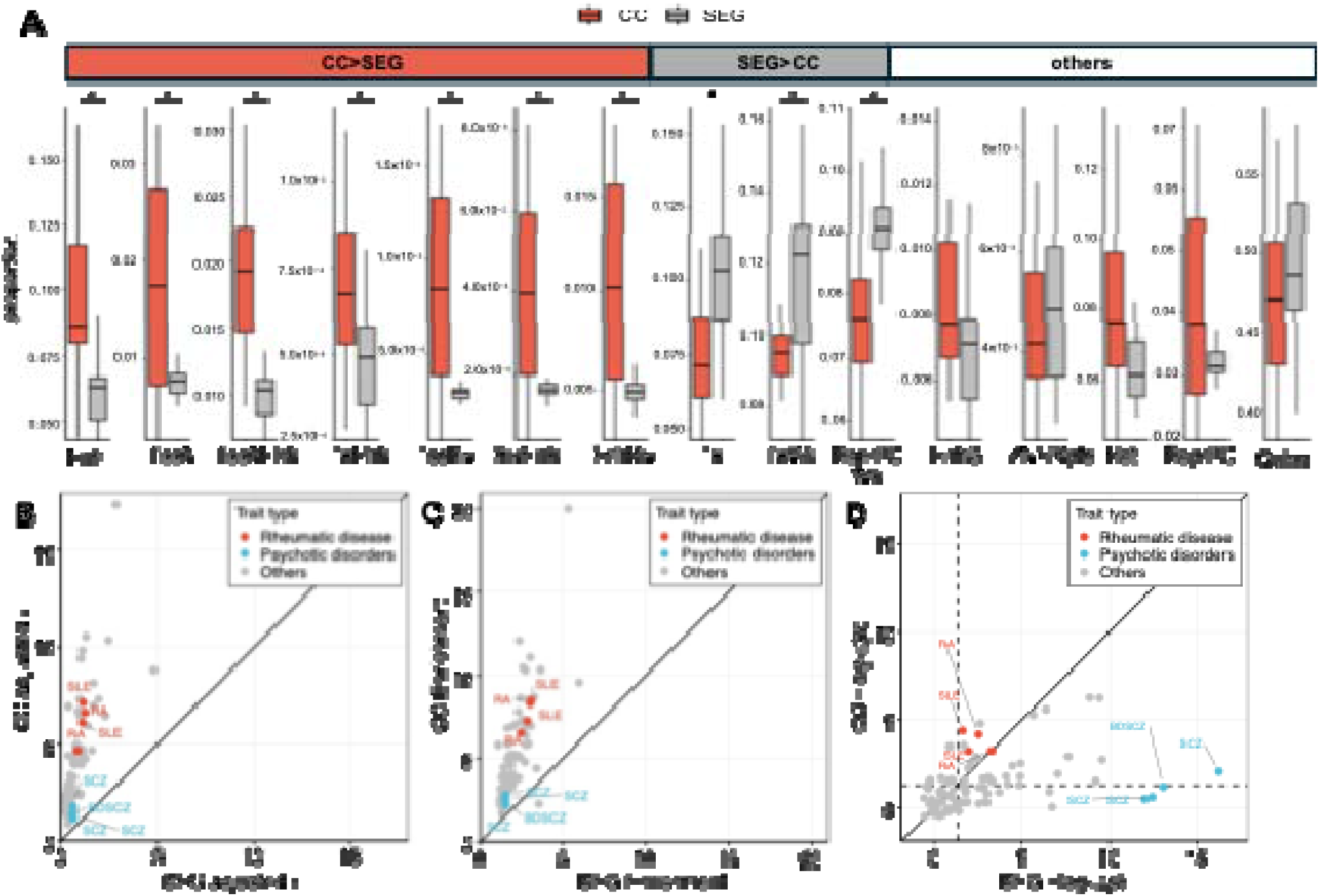
Comparison of CC and SEG annotation tracks. (A) Box plots showing the proportion of SNPs mapping to each chromatin state category defined by the Roadmap Epigenomics core 15-state ChromHMM model (SEG, *n* = 53; CC, *n* = 20). Chromatin state assignments were based on Roadmap sample E029 (primary monocytes from peripheral blood). Asterisks denote categories with significant differences between CC and SEG annotations after Bonferroni correction (Wilcoxon rank-sum test). Chromatin states were defined using the 15-state ChromHMM model. (B–D) Comparison of adjusted *r* (regression coefficient from stratified LD score regression) (B), heritability enrichment (C), and Bonferroni-corrected −log₁₀(*p*) of *r* (D) between CC and SEG annotations (SEG, *n* = 54; CC, *n* = 20, GWAS traits: 89). The black line indicates *y* = *x*. The performance indexes of the annotation track with the highest −log₁₀(*p*) of *r* from each method are shown. Rheumatic diseases are highlighted in red; psychotic disorders in blue. In D, the dashed lines indicate the significance threshold. Enh, Enhancers; TssA, Active TSS; TssAFlnk, Flanking Active TSS; TxFlnk, Transcription at gene 5’ and 3’; TssBiv, Bivalent/Poised TSS; BivFlnk, Flanking Bivalent TSS/Enh; EnhBiv, Bivalent Enhancer; Tx, Strong transcription; TxWk, Weak transcription; ReprPCWk, Weak Repressed PolyComb; EnhG, Genic enhancers; ZNF/Rpts, ZNF genes and repeats; Het, Heterochromatin; ReprPC, Repressed PolyComb; Quies, Quiescent/Low; SLE, systemic lupus erythematosus; RA, rheumatoid arthritis; SCZ, schizophrenia; BDSCZ, bipolar disorder and schizophrenia.

Next, we compared S-LDSC results between CC and SEG annotations. Because CATaN defines multiple orthogonal axes from a single transcriptome, each capturing distinct biological variation, we evaluated the performance of each method by comparing the most significant annotation tracks. CC annotations showed higher adjusted *r* (regression coefficient from stratified LD score regression) and enrichment than SEG annotations across traits (Figures 5B, 5C and Table S5D). This advantage likely arises because CC annotations preferentially capture core regulatory regions, particularly enhancers, where causal variants are known to be concentrated, as shown in Figure 5A. For the significance of heritability enrichment, performance varied by trait: CC annotations tended to outperform SEG for rheumatic diseases (e.g., SLE and RA), whereas SEG tended to outperform CC for psychotic disorders (e.g., schizophrenia) (Figure 5D). This pattern likely reflects the underrepresentation of neural-related TFs in the current TF-GRN matrix, limiting the ability of CC annotations to capture regulatory programs relevant to neuropsychiatric traits. Together, these results indicate that CC and SEG annotations capture partly distinct, complementary aspects of disease heritability.

Beyond trait-dependent differences, the relative performance of CC and SEG annotations also varied by transcriptome dataset type. To illustrate this, we compared the two approaches using RA (EUR) GWAS heritability across two representative transcriptome datasets: GTEx, a multi-tissue dataset, and the multi-autoimmune dataset, a clinical dataset. As above, we evaluated each method by comparing the most significant annotation tracks. CC annotations showed higher enrichment and adjusted *r* than SEG annotations in both datasets (Figures S4C and S4D). However, the two methods diverged in statistical significance. In GTEx, the most significant CC and SEG tracks achieved comparable −log₁₀(*p*) of *r* (CC: Bonferroni-corrected *p* = 6.3 × 10^−4^; SEG: 5.0 × 10^−4^). By contrast, in the multi-autoimmune dataset, the best-performing CC track survived Bonferroni correction (Bonferroni-corrected *p* = 2.0 × 10^−10^), whereas the best-performing SEG track did not (Bonferroni-corrected *p* = 6.3 × 10^−2^) (Figure S4E). This divergence may reflect a difference in how the two methods capture disease-relevant variation in clinical transcriptomes. Pathway analysis revealed that RA DEGs (differentially expressed genes), on which the SEG annotations are based, were enriched for non-inflammatory pathways, whereas genes with high CCA gene projections showed enrichment for inflammatory pathways including viral infection and autoimmune disease pathways (Figure S4F). These results suggest that for clinical transcriptomes, the axis with maximal heritability enrichment does not necessarily align with the disease-versus-control axis captured by SEG. While SEG treats all disease-associated DEGs as a single annotation, CATaN partitions this label-based gene grouping into functionally distinct components through TF-GRN information, potentially isolating the regulatory component most relevant to heritability enrichment.

### SNP-heritability enrichment across diverse transcriptome datasets

We applied S-LDSC to the 740 CC annotation tracks (top and bottom deciles × 10 CCs × 37 datasets) derived from eight biologically diverse transcriptome datasets (Table 1). We quantified SNP-heritability enrichment across 90 traits/diseases. In total, 588 significant enrichments among 63,700 applications were detected, spanning 69 traits and diseases. To confirm cross-ancestry robustness of CC annotation tracks, we compared the adjusted *r* values derived from five bulk datasets between EUR and EAS GWAS for RA and found a high correlation (*r* = 0.77) (Figure S5A).

We next asked how disease-associated heritability distributes across transcriptional heterogeneity. As a representative example, we overlaid the sample loadings of CC annotation tracks significantly enriched for SNP-heritability of type 1 diabetes (T1D) (Figure S5C), major depressive disorder (MDD) (Figure S5D), and prostate and breast cancer (Figure S5E) onto the transcriptome-derived UMAP of GTEx (Figures S5B and Table S5D). Each enriched annotation can be characterized by three CATaN outputs (Tr-loading, TF-loading, and gene projection). The annotation most enriched for MDD heritability (CC1-top) corresponds to brain (Tr-loading), REST and EZH2 (TF-loading), and GABAergic synapse (gene projection); whereas that for T1D (CC2-bottom) corresponds to blood (Tr-loading), PAX5 (TF-loading), and B cell receptor signaling (gene projection).

For other applications, we highlight several notable findings here and report the remaining results in the Supplementary Note and Table S7-15). This framework is also applicable to single-cell data. Applying CATaN to a T cell subset of an RA synovium dataset^28^ (Figure 6A and Table S13), we found that RA heritability was enriched in CC3-bottom, which was enriched for T peripheral helper (Tph) cells (Figure 6B), and in CC5-bottom, which was enriched for GZMK/B⁺ memory CD8⁺ T cells (Figure 6C). In the B cell subset (Figure 6D and Table S14), RA heritability was enriched in CC1-bottom, which was enriched for germinal center-like (GC-like) B cells (Figure 6E), suggesting a role for ectopic germinal center formation in RA, and in CC2-top, which was enriched for autoimmune-associated B cells (ABCs) ^29^(Figure 6F). While these cell populations have been implicated in RA pathogenesis^30^, heritability enrichment had previously been reported for CD4⁺ T cell states, in particular Tph cells^13^. CATaN additionally detected enrichment in B cell populations, namely GC-like B cells and ABCs. In neural organoid data^31^ (Figure 6G and Table S15), major depressive disorder (MDD) showed the greatest heritability enrichment in CC2-top, which was enriched for radial glial cells and mature neurons (Fig 6H). Bipolar disorder exhibited the greatest enrichment in CC3-bottom, which was enriched for neuronal cells (Figures 6G). MDD has long been attributed to abnormalities in monoaminergic transmission in neuronal cells; however, recent snRNA-seq studies^32^ of the brain have reported that differentially expressed genes of MDD are concentrated not only in neurons, but also in immature oligodendrocyte precursor cells (OPCs). Radial glial cells are multipotent progenitors that can give rise to the oligodendrocyte lineage as well as to neurons and astrocytes^33^. Our finding of heritability enrichment in these cells is therefore consistent with recent reports implicating oligodendrocyte-lineage cells, including immature OPCs, in the pathophysiology of MDD^32^.

**Fig. 6.**
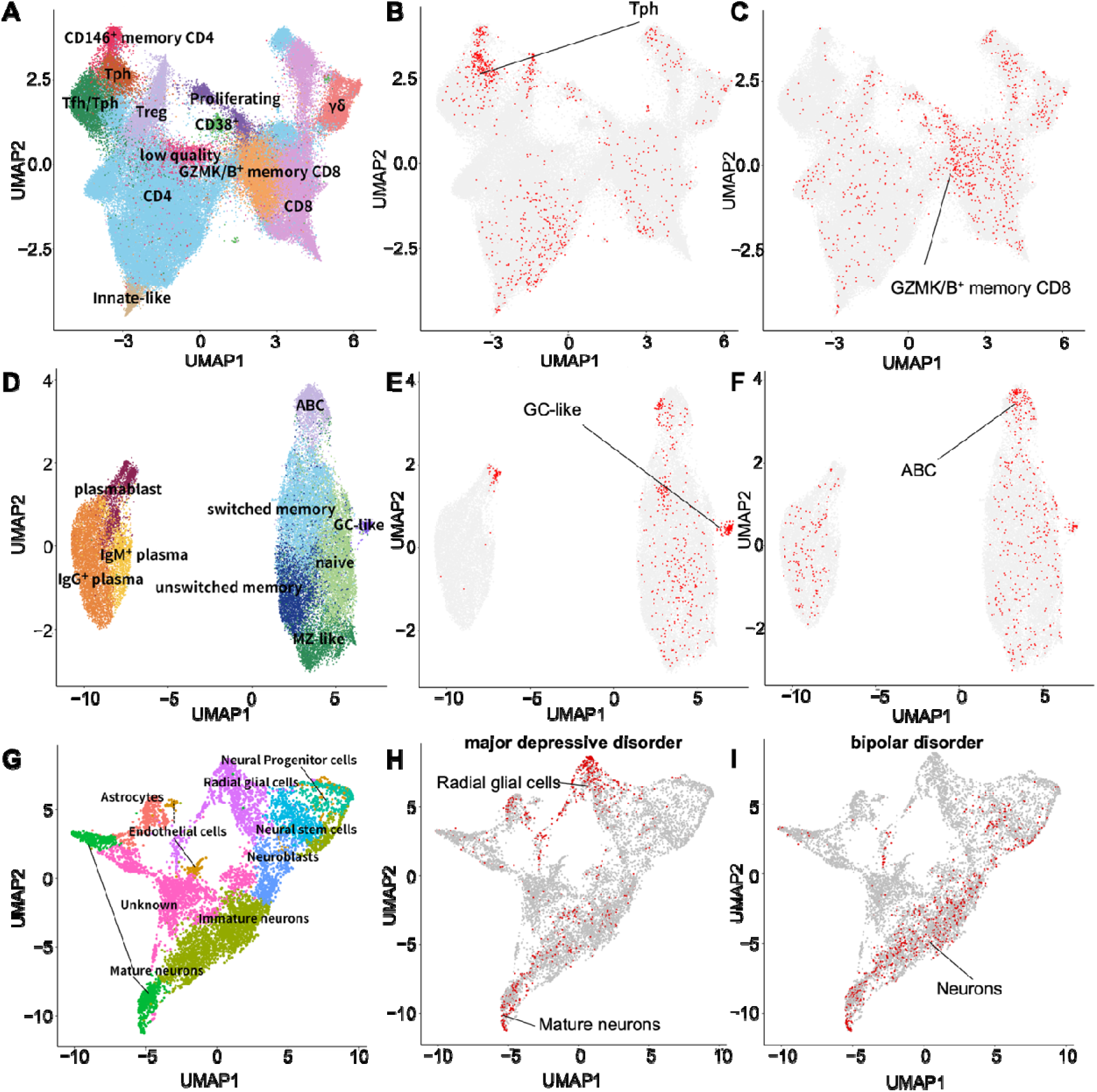
Heritability enrichment mapped onto transcriptome UMAP space. (A–F) RA synovium dataset. (A) UMAP of T cell sub-dataset (*n* = 94,046) colored by cell types. (B–C)Top or bottom 2% of Tr sample loadings from CC annotation tracks enriched for RA heritability, highlighted in red (B, CC3 bottom; C CC5 bottom). (D) UMAP of B cell sub-dataset (*n* = 30,691) colored by cell types. (E–F) Top or bottom 2% of Tr sample loadings from the CC annotation tracks enriched for RA heritability, highlighted in red (E, CC1 bottom; F, CC2 top). (G–I) Neural organoid dataset (*n* = 8,295). (G) UMAP colored by scType cell-ype annotation. (H) Top or bottom 10% of Tr sample loadings from the most heritability-enriched CC for major depressive disorder (MDD), highlighted in red. (I) As in H, for bipolar disorder. GC-like, germinal center-like B cells. ABC, autoimmune-associated B cells. Tph, T peripheral helper cells.

### Proof-of-principle functional variant prioritization using CATaN

Because CC annotation tracks show broad SNP-heritability enrichment and provide single-base-p air resolution independently of linkage disequilibrium (LD), we hypothesized that per-SNP CC scores could help prioritize candidate causal variants.

We used RA, a trait with a large number of fine-mapped GWAS variants, to assess whether CC annotations are enriched for fine-mapped variants. We observed that the CC track with the highest S-LDSC significance for RA heritability enrichment also showed the greatest enrichment of fine-mapped RA GWAS variants^34^ (Figure 7A). We then experimentally compared caQTL effects between GWAS lead SNPs located outside CC annotations and their LD partners located within CC annotations (CC SNPs), selected on the basis of functional evidence from massively parallel reporter assays (MPRA)^35^ (Figure 7B; see Methods). Of the three tested regions, only one, a T1D-associated locus, achieved sufficient editing efficiency for analysis (Figure S6), and we therefore used this locus for caQTL validation as a proof of principle. Using UNIChro-seq^36^, a targeted chromatin accessibility assay developed in our laboratory, the CC SNP rs74548327 (chr17:45,895,714:A>G, hg38) (effect size: 0.45, *p* = 8.9 × 10^−6^) exhibited a larger caQTL effect than the corresponding T1D GWAS^37,38^ lead SNP rs1052553 (chr17:45,996,523:A>G, hg38) (effect size: −0.21, *p* = 7.6 × 10^−2^) (Figure 7C, Table S17). Although limited in scale, these results provide proof-of-principle evidence that CATaN-derived scores may aid in the prioritization of functional variants for experimental follow-up.

**Fig. 7.**
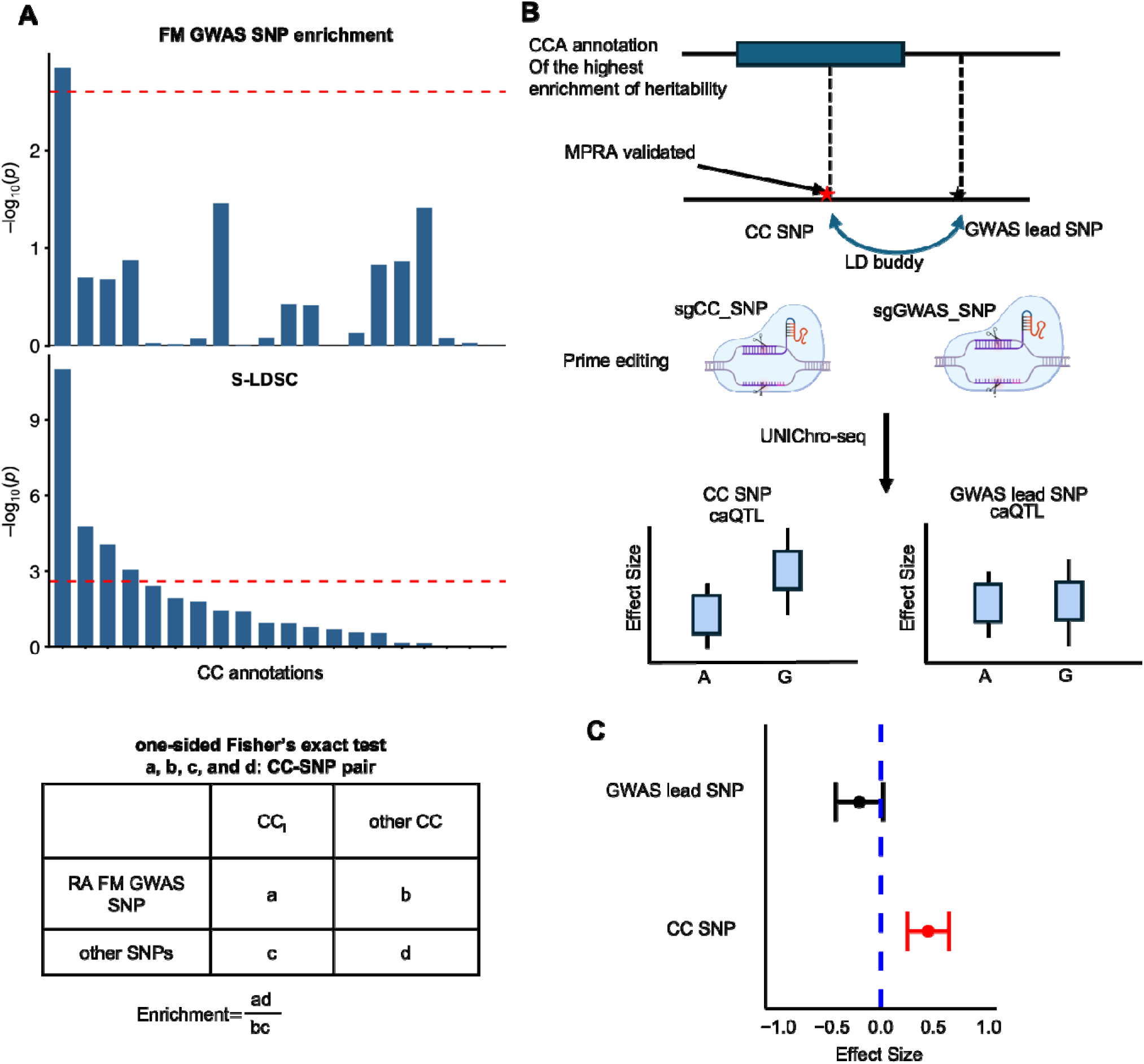
Proof-of-principle variant prioritization using CATaN. (B) Enrichment of RA GWAS heritability and fine-mapped (FM) variants in CC annotations. The result of double-negative B cells in the multi-autoimmune dataset is shown as representative. Top: one-sided Fisher’s exact test was used to assess enrichment of RA FM GWAS variants in each CC annotation. A 2 × 2 contingency table (bottom) was constructed, classifying each CC–SNP pair by fine-mapped GWAS variant status (posterior inclusion probability (PIP) > 0.2) and inclusion in a given CC annotation versus other CC annotations. Middle: the x axis shows 20 CC annotations ranked by S-LDSC heritability enrichment significance in descending order. Dashed lines indicate the Bonferroni-corrected significance threshold. Top, −log₁₀(*p*) of fine-mapped variant enrichment (one-sided Fisher’s exact test); bottom, −log₁₀(*p*) of *r* from S-LDSC. (C) Schematic illustrating experimental fine-mapping. caQTL effects between GWAS lead SNPs (GWAS SNPs) located outside CC annotations and their LD partners located within CC annotations (CC SNPs). We selected CC SNPs with functional evidence from massively parallel reporter assays (MPRA). CC SNPs and GWAS SNPs were edited using prime editing, and caQTLs were measured using UNIChro-seq. (D) Comparison of chromatin accessibility QTL (caQTL) effect sizes measured by UNIChro-seq between a CC SNP and the corresponding type 1 diabetes (T1D) GWAS lead SNP. GWAS lead SNP is rs1052553 (chr17:45,996,523:A>G, hg38). CC SNP is rs74548327 (chr17:45,895,714:A>G, hg38).

## Discussion

In this study, we developed CATaN, an unsupervised framework that integrates transcriptome data with TF-mediated gene regulatory networks via CCA to identify genome-wide annotation tracks enriched for disease heritability. CATaN differs from conventional approaches in three major ways. First, it is fully unsupervised, enabling comprehensive exploration of transcriptomic variance without relying on predefined labels, an advantage when axes of transcriptomic variation and heritability enrichment are not aligned. Second, beyond CC annotation tracks, CATaN provides TF sample loadings, transcriptome sample loadings, and gene projections, facilitating mechanistic interpretation in terms of specific TFs, sample metadata, and gene programs. Third, CC annotation tracks are built from TF sample loadings combined with ChIP-seq binding profiles, in contrast to proximity to gene bodies alone. Collectively, these features make CC tracks a powerful resource for quantifying heritability enrichment while simultaneously providing mechanistic insight and facilitating the prioritization of candidate causal variants.

CATaN can identify the specific cell types and biological states enriched for disease heritability without relying on predefined labels. In the T cell and B cell subsets of the RA synovium dataset, heritability-enriched cell types were identified. These findings are consistent with a previous multi-omics study integrating scATAC-seq and scRNA-seq^13^, which reported heritability enrichment in overlapping T cell populations. Notably, CATaN recovered these T cell associations using scRNA-seq data alone, highlighting its broader applicability to datasets where chromatin accessibility data are unavailable. Furthermore, CATaN additionally detected enrichment in B cell populations, including ABCs, which are implicated in RA pathogenesis^39^, and GC-like B cells associated with ectopic germinal center formation^40^, but had not been highlighted in heritability analyses based on chromatin accessibility^13^. Moreover, while the previous multi-omics study did not detect heritability enrichment in GZMK/B⁺ memory CD8⁺ T cells and proposed that their expansion reflects a secondary response to inflammation^13^, CATaN identified enrichment in this population. This difference may arise because CATaN captures TF-GRN-based regulatory signal rather than chromatin accessibility, although the contribution of GZMK/B⁺ CD8⁺ T cells to RA heritability remains to be established. More broadly, methods to project disease heritability enrichment onto single-cell transcriptomic landscapes in a label-free manner remain scarce. Unlike scDRS, which integrates GWAS data at the gene level, CATaN operates at the SNP level, retaining genetic information that is aggregated in gene-based approaches such as scDRS. CATaN addresses this gap in label-free methodology by providing a systematic framework to localize heritability signals within cellular heterogeneity, and we anticipate that its application to diverse single-cell transcriptome datasets will yield further biological insights.

Our study presents four primary limitations. First, the TF ChIP-seq dataset was assembled by broadly curating all available public data at the time of analysis; nevertheless, substantial biases exist in both antibody targets and sampled tissues. For example, CTCF accounts for approximately 10% of all TFs, and immune-related TFs such as RELA and STAT family members are heavily represented. Likewise, approximately one-third of samples originate from immune cells. These imbalances reflect the current constraints in the quantity of publicly available ChIP-seq resources. Expansion of ChIP-seq datasets, particularly across diverse TFs and tissue contexts, will help mitigate these biases and is expected to further improve method performance.

Second, potential inaccuracies exist in model specification, including the strategy used to link TF binding sites to genes. At present, no experimental gold standard exists for validating TF-gene regulatory links in a genome-wide manner, making this an important area for future methodological and experimental development. Third, our experimental fine-mapping analysis was limited to a single locus where both the GWAS lead variant and the CC SNP achieved sufficient editing efficiency for caQTL comparison. Low editing efficiency, particularly at GWAS lead variants, was the primary bottleneck, possibly reflecting reduced chromatin accessibility at these sites. Expanding the experimental validation to additional loci will be necessary to more robustly evaluate the utility of CC annotation for causal variant prioritization. Fourth, our single-cell heritability enrichment analyses were performed at the cell level, where cells from the same donor are not statistically independent. The cell-type-level signals we report therefore require validation with donor-aware approaches in larger cohorts.

CATaN is a versatile framework applicable across diverse transcriptomic datasets. By structuring transcriptomic variance through the lens of TF-GRN information, it identifies components associated with disease heritability and paves the way for prioritizing regulatory programs and candidate variants in basic and clinical research.

## Methods

### TF ChIP-seq data curating method

The processing was performed as previously described^6,41^. Briefly, we downloaded 3,158 human ChIP-seq datasets from the Gene Expression Omnibus (GEO) database. We mapped these reads to the genome assembly GRCh37 using Bowtie 2^42^ (version 2.2.5) with default parameters. We called peaks using MACS^43^ (version 2.1) with default parameters (q < 0.01) and defined them as TF binding sites. We excluded TF binding site tracks that did not have at least one binding region on every chromosome, and 2,868 genome-wide TF binding site tracks remained.

### TF-GRN matrix

A TF-GRN matrix (rows: genes, columns: TF ChIP-seq samples) was constructed based on our curated TF ChIP-seq dataset, encompassing 2,868 samples representing 410 distinct TFs. To establish gene-TF associations, we adopted the exponential distance-based weighting scheme implemented in MAESTRO^44^. Each TF peak is weighted by an exponential function of its distance from the transcription start site (TSS), with these weighted values subsequently aggregated (half-decay distance: 10 kbp) as equation (1):

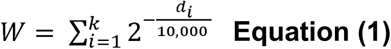

Where *w* corresponds to TF-gene connectivity scores, *k* corresponds to the number of ChIP-seq peaks within 1 Mbp of the TSS of each gene, and *d_i_* corresponds to the distance between peak i and the TSS of the gene. For TSS annotation data, we utilized GENCODE Release 26 (GRCh37). We evaluated the validity of the weighting parameters using singular values (SVs). The SVs serve as an index of how well CCA extracts the shared variation between the TF-GRN matrix and the transcriptome matrix, with larger SVs indicating stronger shared components. For each weighting condition, we summed the first ten SVs (SV1–SV10), matching the number of CC components used in the downstream analyses, and compared this sum across conditions. Exponential weighting with a 10 kb half-decay distance yielded the highest summed SVs, outperforming a binary approach and other half-decay distances (Supplementary Note,Figures S1B and S1C); this weighting was therefore used for all analyses in this study.

### Transcriptome data matrix

For RNA-seq data, we retained genes with more than 10 counts and more than 1 CPM, each in more than 15% of samples, following a previously described filtering approach^45^, using the pOverA filter in the genefilter package^46^. We then retained the intersection of genes with TF-GRN matrix. Counts were normalized by a trimmed mean of M values (TMM) normalization with edgeR (version 4.0.16)^47^. Normalized expression data were converted to log-transformed CPM (log_2_(CPM+1)). This transformation resulted in a transcriptome data matrix with genes represented in rows and samples in columns. For microarray data analysis, we utilized the phase 1 dataset (GSE15907) obtained from the Immunological Genome Project (ImmGen) Consortium. The data, originally uploaded to the GEO database in exponential scale, were subjected to log_2_ transformation. Subsequently, we converted the mouse genes to their human orthologs using ENSEMBL orthologs database. For single-cell RNA-seq datasets, we used the processed Seurat objects provided by the original studies. We normalized the raw count data using log-transformation:

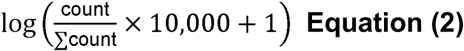

equivalent to Seurat’s LogNormalize with a scale factor of 10,000. We included genes expressed in at least 1% of cells. For cluster annotation, we applied scType^48^ to the neural organoid dataset and Azimuth^49^ to the ILC dataset^50^; for the RA synovium dataset, we used the cell-type annotations provided by the original study.

### CCA

Because the large number of samples made the covariance matrix computation intractable for conventional CCA, we used the diagonal CCA approach implemented in Seurat^51^ as described in equation (3).

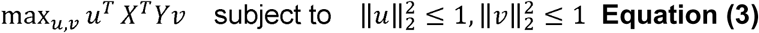

where *X* and *Y* correspond to scaled TF-GRN matrix and transcriptome matrix, *u* and *v* correspond to the TF sample loadings and transcriptome sample loadings.

To provide a more detailed explanation, we first scale each gene to mean = 0, sd = 1 (gene-wise), and then each sample (sample-wise) as equations (4) and (5).

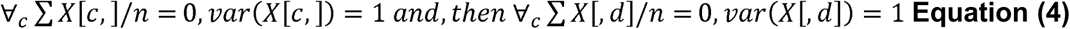

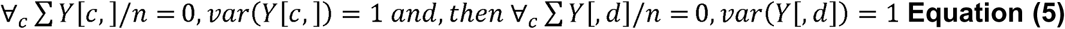

where *X* corresponds TF-GRN data matrix and *Y* corresponds transcriptome matrix.

The top 10,000 most variable genes were extracted from both the TF-GRN data matrix and the transcriptome matrix. We limited genes to the intersection of these two most variable gene populations. Subsequently, we performed singular value decomposition (SVD) to obtain the canonical correlation (CC) basis scores as follows: Let

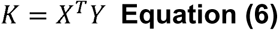

K can be decomposed using SVD as

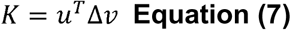

where *Δ* denotes singular values (SVs), 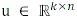 denotes TF sample loading, and 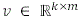 denotes transcriptome sample loading, with k denoting the number of canonical components, n the number of TF ChIP-seq samples, and m the number of transcriptome samples. Gene projections were obtained by multiplying the pre-CCA matrices with their respective sample loading matrices. Two types of gene projections were generated: TF gene projection and transcriptome gene projection. For subsequent analyses, we used only the transcriptome gene projection due to redundancy (Supplementary Note, Figure S1D). Transcriptome sample loading, TF sample loading and gene projection were converted to Z-scores by subtracting the mean and dividing by the standard deviation across all samples/genes when referenced in the text. For the multi-autoimmune dataset, which includes control samples, the mean and standard deviation of the control samples were used instead.

### Enrichment analysis of sample loadings

To evaluate whether CATaN sample loadings are enriched for specific metadata labels, we performed one-sided Fisher’s exact test. For each CC axis, we defined the top-ranked and bottom-ranked samples based on Z-scored transcriptome sample loading and tested each set separately against the remaining samples using one-sided Fisher’s exact test. The rank threshold was set at 500 in GTEx, at 300 in ImmGen, at 80 in human immune cell dataset and single-cell datasets, at 60 in the multi-autoimmune dataset, and at 20 in cytokine CD4 dataset for transcriptome sample loadings, and at 200 for Z-scored TF sample loadings. Metadata labels were defined according to each dataset (e.g., tissue type, cell type, or disease status for transcriptome samples; target TF for ChIP-seq samples). P values were corrected for multiple testing using the Bonferroni method.

### Index for the efficacy of CCA

The SV reflect the covariance components shared between the transcriptome and TF-GRN matrices. This value serves as an indicator of the efficacy of our CCA model. To rigorously assess the statistical significance of our results, we implemented a null model, created by permuting the genes in the transcriptome data matrix (Figure S3A). This process effectively destroys the data structure while preserving the overall variance of the transcriptome matrix and TF-GRN matrix. The permutation was conducted 1000 times. We calculated the ratio of the observed SVs to the mean values from the null model as follows:

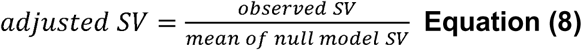

### Variance explained by TF and transcriptome sample loading

We defined the variance explained by each canonical component (CC) axis as follows.

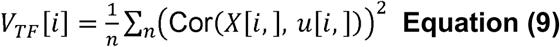

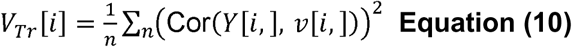

where *v_TF_*: TF sample loading, *v_Tr_*: variance explained by transcriptome sample loading, *i*: the order of CC, *n*: the number of genes in *X* and *Y*.

### Variance partitioning of transcriptome and TF sample loading using metadata

The variance partitioning of transcriptome and TF sample loadings was inspired by variance partitioning in PCA. For TF sample loadings, we fitted the following generalized linear model:

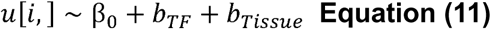

where *β*_0_ is the intercept, *b_TF_* is a random effect for TF identity, and *b_Tissue_* is a random effect for tissue. We then calculated the variance attributed to each random effect term and residual term. Then, we computed their relative proportions. For transcriptome sample loadings, we applied the same procedure to the following model:

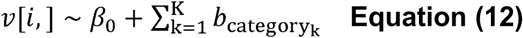

where *β*_0_ is the intercept and *b_Categoryk_* is a random effect for the *k*-th category.

### Creation of CC annotation tracks from TF sample loadings

We constructed annotations for S-LDSC using TF sample loadings derived from CCA. The process involved the following steps (Figure 1B):

1. **TF Peak Region Scoring**: We assigned TF sample loadings to TF peak regions.
2. **SNP-level Score Aggregation**: For each SNP used in LD score calculation, we summed the TF sample loadings of all TF peak regions overlapping that SNP. SNPs not located within any TF binding site were assigned a score of 0.
3. **Binary Annotation Creation**: The top 10% and bottom 10% of SNPs by score were each used to define a separate binary annotation.
4. We applied S-LDSC using the baseline-LD model (version 1.2) in conjunction with our custom CC annotations. Analyses were restricted to HapMap3 SNPs^52^, excluding the MHC region. Pre-computed LD scores, regression weights, and allele frequencies were obtained from the Alkes Group repository. The 1000 Genomes Phase 3 European reference panel was used for European-ancestry GWAS, and the corresponding East Asian reference panel for East Asian-ancestry GWAS. We obtained European-ancestry GWAS summary statistics for 89 traits and diseases from the repository maintained by the Price laboratory (https://alkesgroup.broadinstitute.org/sumstats_formatted/). Sources include summary statistics curated by Finucane et al.^9^, UK Biobank^53^, and individual GWAS consortia (Table S16). For cross-ancestry analyses, East Asian-ancestry GWAS summary statistics were obtained for RA from Ishigaki et al.^34^. We assessed both the effect size and statistical significance of heritability enrichment for each annotation, using adjusted *r* and p value of *r*, where

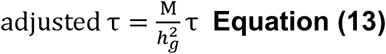

M: the number of SNPs used to compute 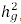, which is a corrected for the number of SNPs used in LD score calculation and total SNP heritability, inspired by the previous paper^54^. M is 5,926,598 for European GWAS summary statistics, and 5,633,366 for East Asian GWAS summary statistics.

### Pathway Analysis of gene projection

We conducted an Over-Representation Analysis (ORA) to examine the enrichment of pathway genes from MSigDB and KEGG databases within the top and bottom 10% of genes in the transcriptome gene projection. The clusterProfiler R package (version 4.10.1) ^55^ was employed for the analysis.

### Validation of SNP CC scores using fine-mapped eQTL results

We utilized PIP>0.9 fine-mapping cis-eQTL data from GTEx (version 10) analyzed using SuSiE^56^ (See **Datasets**). We computed eQTL-gene CC scores by averaging CC scores of eVariants per eGENE, generating a gene-length vector (Figure 3E). The resulting scores were then normalized via inverse normal transformation (INT). We assessed its Pearson correlation with Z-scored gene projections.

### Holdout validation using GTEx transcriptome data

GTEx samples were randomly divided into two equal subgroups (training and held-out). CATaN was applied to the training subgroup to derive transcriptome gene projection (gene × CC1-10 matrix). Transcriptome matrices of training and holdout subgroups were independently in the same manner as written in CCA paragraph (first gene-wise, then sample-wise). Transcriptome gene projection of training data was used as loadings to project both training and holdout samples onto the same CC space (Figure 3C).

### CRISPRa-mediated RELA activation and RamDA-seq analysis

Jurkat cells stably expressing dCas9-VP64 were generated by lentiviral transduction with pZR112_Lenti-SFFV-mCherry-2A-dCas9-VP64 (Addgene #180263), followed by fluorescence-activated cell sorting of mCherry-positive cells. For CRISPRa-mediated activation of RELA, the stable Jurkat cells were transduced with pooled lentiviral vectors based on pXPR_502 (Addgene #96923) expressing three distinct single-guide RNAs (sgRNAs) targeting RELA (sgRELA) or a single control sgRNA (sgControl). Following transduction, cells were selected with puromycin at 2 μg/mL for 3 days. The sgRNA sequences are provided in Table S18. Transductions were performed in triplicate for each condition (n = 3). Cells were harvested at 5, 6, and 7 days after transduction and prepared in duplicate for RamDA-seq analysis^57^. All samples were used for differentially expressed gene (DEG) analysis. For CATaN analysis, only day 7 samples were used due to substantial batch effects between time points that could not be adequately corrected. Because this dataset contained fewer samples than the others, the low-expression gene filter was set to retain genes with more than 10 counts and more than 0.5 CPM in more than 5% of samples. DEG analysis was performed using the edgeR package. Gene set enrichment analysis (GSEA) was performed on a ranked gene list using the clusterProfiler package with MSigDB and KEGG databases.

### Enrichment of tissue SEGs and disease DEGs near related TF ChIP-seq peaks in the TF–GRN matrix

Tissue SEGs were defined using GTEx V6p data by comparing expression levels of each tissue against all other tissues using edgeR. SLE DEGs were obtained from a previously reported disease-state differential expression analysis^45^. Tissue-specific TFs were as defined by the GTEx Consortium^58^. Only tissues in which at least 30 tissue-specific TFs were present in the TF–GRN matrix were included in the analysis. A gene was considered proximal to a TF binding site if the corresponding TF–GRN matrix score was ≥1. Enrichment of tissue up– or down-regulated SEGs (or disease up– or down-regulated DEGs) near binding sites of tissue-specific TFs (or disease-related TFs) or other TFs was tested using a one-sided Fisher’s exact test.

### Chromatin state enrichment analysis

To compare chromatin state distributions between SEG and CC annotations, we used the Roadmap Epigenomics core 15-state ChromHMM model. Chromatin state segmentation data for E029 (primary monocytes from peripheral blood) and E071 (Brain Hippocampus Middle) were downloaded from the Roadmap Epigenomics Project (See **Datasets**). For each SNP included in the SEG annotations (*n* = 53) and CC annotations (*n* = 20), we determined the corresponding chromatin state assignment based on the E029 and E071 15-state model. We then calculated the proportion of SNPs mapping to each chromatin state category relative to the total number of SNPs within each annotation. Differences in chromatin state proportions between SEG and CC annotations were assessed using the Wilcoxon rank-sum test.

### Enrichment of RA GWAS fine-mapped variants in CC annotation

We used the double-negative B cell subset of the multi-autoimmune dataset as representative data, as it contained the CC annotation with the highest significance for RA heritability enrichment among all CC annotations analyzed in this study. Fine-mapped (FM) variants from RA GWAS with a posterior inclusion probability (PIP) > 0.2 were used. For each CC annotation, a one-sided Fisher’s exact test was performed to assess the enrichment of fine-mapped GWAS variants, by classifying each CC-SNP pair according to whether the variant was a FM GWAS variant (PIP > 0.2) and whether it was included in the given CC annotation versus other CC annotations.

### Comparison of caQTL effects between GWAS lead variants and CC annotation-included variants

We performed prime editing in Jurkat cells to introduce nucleotide substitutions at eight genomic variants, including five autoimmune disease GWAS lead variants located outside CC annotations and three high-LD variants located within CC annotations showing the strongest heritability enrichment for the corresponding trait. CC SNPs were selected based on functional evidence from massively parallel reporter assays (MPRA). Sites heterozygous in Jurkat cells were selected for analysis, and bidirectional editing was performed separately for REF-to-ALT and ALT-to-REF conversion. Jurkat cells were electroporated with pCMV-PEmax-PE2-GFP (Addgene #180020), pEF1a-hMLH1dn (Addgene #174824), epegRNA expression plasmids cloned into pU6-tevopreq1-GG-acceptor (Addgene #174038), and nicking sgRNA expression plasmids cloned into pFYF1548 EMX1 (Addgene #47508), as previously described^36^. GFP-positive cells were sorted 1 day after electroporation and cultured for 14 days. Eight biological replicates were analyzed for each editing condition. Editing efficiency was quantified by targeted amplicon sequencing. For chromatin accessibility analysis, 20,000 cells per replicate were subjected to UNIChro-seq as previously described^36^. Editing efficiency was calculated using CRISPResso2^59^, and only GWAS lead SNP–CC SNP pairs for which both variants achieved an editing efficiency of at least 25% were included in downstream caQTL analyses (Figure S6). Sequences of the epegRNAs, nicking sgRNAs, primers, and in silico probes used for genome editing and downstream analyses are provided in Table S19 and S20.This resulted in one analyzable region (one GWAS SNP-CC SNP pair). CaQTL effects were evaluated using UNIChro-seq, developed in our laboratory, following previously described methods. Briefly, lme4 was used for regression with the following formula:

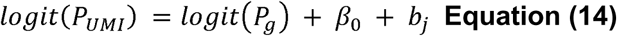

where *P_UMI_* indicates the ALT allele ratio calculated by UMI of UNIChro-seq, *P_g_* is an offset term indicating the ALT allele ratio calculated by reads count of genome DNA-seq. *β_0_* is a fixed effect intercept term indicating the effect size estimate on the chromatin accessibility, and *b_j_* is a random intercept term for the *j*-th replicate sample.

### Cytokine CD4 dataset

The study protocol was approved by the Ethics Committees of the University of Tokyo (approval number G3582). Written informed consent was obtained from all participants. Naïve CD4⁺ T cells were isolated from 28 healthy donors and cultured under five polarization conditions (Th0, Th1, Th2, Th17, and iTreg), yielding 138 samples in total (Table S21). Peripheral blood mononuclear cells (PBMCs) were isolated via density gradient centrifugation using Ficoll-Paque Plus (GE Healthcare). CD3^+^CD4^+^CD45RA^+^CD25^-^ naïve T cells were subsequently sorted by fluorescence-activated cell sorting (FACS) using MoFlo XDP (Beckman Coulter). Antibodies used for sorting are detailed in Table S22. Isolated cells were cultured in RPMI 1640 medium (Invitrogen) supplemented with 10% fetal calf serum (FCS; Equitech Bio), 100 μg/ml L-glutamine, 100 U/ml penicillin, 100 μg/ml streptomycin (all from Invitrogen), and 50 μM 2-ME (Sigma) at 5×10^4^ cells/well. For Th0, Th1, Th2, Th17, and iTreg polarization, cells were cultured in 96 well round-bottom plates that had been precoated with 10μg/ml anti-CD3 antibody (R&D, MAB100, clone UCHT1) overnight at 4℃. All culture conditions were performed in duplicate. Cytokines and antibodies added for each polarization condition are listed in Table S23. After 72 hours of culture, cells were lysed with RLT buffer (QIAGEN). Total RNA was extracted using AllPrep DNA/RNA/miRNA Universal Kits (QIAGEN). RNA-seq libraries were prepared using the TruSeq Stranded mRNA Library Prep Kit (Illumina), and sequencing was performed on an Illumina HiSeq 2500 platform.

### Datasets

*GTEx.* We downloaded RNA-seq read counts from GTEx v6p (https://gtexportal.org/). We used the “SMTS” variable to define tissue annotations. Fine-mapping cis-eQTL data analyzed with SuSiE were obtained from GTEx v10 (https://www.gtexportal.org/home/downloads/adult-gtex/qtl).

*ImmGen.* We downloaded publicly available microarray gene expression data on 653 samples from the ImmGen Consortium (phase 1; GEO accession: GSE15907). Data were log₂-transformed and mapped to human orthologs using the ENSEMBL orthologs database.

*ImmuNexUT.* We downloaded publicly available RNA-seq read count data from the NBDC Human Database (accession: E-GEAD-397).

*Single-cell RNA-seq datasets.* We downloaded the neural organoid dataset (GEO: GSE163018), mouse ILC dataset (GEO: GSE149622), and RA synovium dataset (AMP Phase II) (Synapse: syn52297840).

Roadmap Core 15-state model. We downloaded the core 15– state model from Roadmap Epigenomics Project: https://egg2.wustl.edu/roadmap/data/byFileType/chromhmmSegmentations/ChmmModels/coreMarks/jointModel/final/

## Data availability

CATaN source code and R package are available at https://github.com/htakahashi1/CATaN. Processed data are available at Zenodo (https://doi.org/10.5281/zenodo.19507630). RamDA-seq analysis of CRISPRa-mediated RELA activation is available at GSE335951.

## Acknowledge

This work was supported by JSPS Grant-in-Aid (JSPS Fellows (24KJ1971); Scientific Research (B) (22H03114); Research Activity Start-up (21K20647); and Early-Career Scientists (25K19621)), the Japan Agency for Medical Research and Development (AMED) (JP22tm0424223; JP22ek0410099; JP23tm0524005; JP25ek0410139; JP22ama121015; and JP223fa627010), Keio University Academic Development Funds, Program for the Advancement of Next Generation Research Projects, GSK Japan Research Grant 2021, Daiichi Sankyo Foundation of Life Science, Mochida Memorial Foundation for Medical and Pharmaceutical Research, The Uehara Memorial Foundation, Takeda COCKPI-T Funding, and funding from Takeda Pharmaceutical (Y.K., K.Y. and K.F.). The super-computing resource was provided by Human Genome Center, Institute of Medical Sciences, The University of Tokyo and RIKEN. We appreciate the RIKEN-IMS Genome Platform for their help in the sequencing experiment for this study. We also thank Eiryo Kawakami and Hiroki Sugishita for kindly sharing their results of ChIP-seq data analysis.

## Author contributions

H.T., H.H., and K.I. conceived the project; H.T., H.H., and K.I. wrote the manuscript with support from M.K., M.N., H.I., K.A., T.N., Y.K., T.I., J.I., B.N. and Y.T.; H.T. developed the CATaN pipeline with substantial support from K.I and H.H.; K.H., M.M.D., Y.O. and D.H. analyzed single cell transcriptome datasets and supported CATaN’s performance evaluation; M.K. R.B, T.K., T.A. and B.N. conducted functional experiments using genome editing, UNIChro-seq and CRISPRa; Y.T., S.S, A.S., Y.K., K.F., and K.Y. conducted experiment of cytokine CD4 dataset; T.O. and E.K. analyzed TF ChIP-seq data.

## Reference

1. Laufer, V.A., Tiwari, H.K., Reynolds, R.J., Danila, M.I., Wang, J., Edberg, J.C., Kimberly, R.P., Kottyan, L.C., Harley, J.B., Mikuls, T.R., et al. (2019). Genetic influences on susceptibility to rheumatoid arthritis in African-Americans. Hum. Mol. Genet. 28, 858–874. 10.1093/hmg/ddy395.

2. Okada, Y., Wu, D., Trynka, G., Raj, T., Terao, C., Ikari, K., Kochi, Y., Ohmura, K., Suzuki, A., Yoshida, S., et al. (2014). Genetics of rheumatoid arthritis contributes to biology and drug discovery. Nature 506, 376–381. 10.1038/nature12873.

3. O’Connor, L.J., Schoech, A.P., Hormozdiari, F., Gazal, S., Patterson, N., and Price, A.L. (2019). Extreme Polygenicity of Complex Traits Is Explained by Negative Selection. Am. J. Hum. Genet. 105, 456–476. 10.1016/j.ajhg.2019.07.003.

4. Finucane, H.K., Bulik-Sullivan, B., Gusev, A., Trynka, G., Reshef, Y., Loh, P.-R., Anttila, V., Xu, H., Zang, C., Farh, K., et al. (2015). Partitioning heritability by functional annotation using genome-wide association summary statistics. Nat. Genet. 47, 1228–1235. 10.1038/ng.3404.

5. Johnson, D.S., Mortazavi, A., Myers, R.M., and Wold, B. (2007). Genome-wide mapping of in vivo protein-DNA interactions. Science 316, 1497–1502. 10.1126/science.1141319.

6. Ishigaki, K., Akiyama, M., Kanai, M., Takahashi, A., Kawakami, E., Sugishita, H., Sakaue, S., Matoba, N., Low, S.-K., Okada, Y., et al. (2020). Large-scale genome-wide association study in a Japanese population identifies novel susceptibility loci across different diseases. Nat. Genet. 52, 669–679. 10.1038/s41588-020-0640-3.

7. Amariuta, T., Luo, Y., Gazal, S., Davenport, E.E., van de Geijn, B., Ishigaki, K., Westra, H.-J., Teslovich, N., Okada, Y., Yamamoto, K., et al. (2019). IMPACT: Genomic Annotation of Cell-State-Specific Regulatory Elements Inferred from the Epigenome of Bound Transcription Factors. Am. J. Hum. Genet. 104, 879–895. 10.1016/j.ajhg.2019.03.012.

8. Martin-Rufino, J.D., Caulier, A., Lee, S., Castano, N., King, E., Joubran, S., Jones, M., Goldman, S.R., Arora, U.P., Wahlster, L., et al. (2025). Transcription factor networks disproportionately enrich for heritability of blood cell phenotypes. Science 388, 52–59. 10.1126/science.ads7951.

9. Finucane, H.K., Reshef, Y.A., Anttila, V., Slowikowski, K., Gusev, A., Byrnes, A., Gazal, S., Loh, P.-R., Lareau, C., Shoresh, N., et al. (2018). Heritability enrichment of specifically expressed genes identifies disease-relevant tissues and cell types. Nat. Genet. 50, 621–629. 10.1038/s41588-018-0081-4.

10. Jagadeesh, K.A., Dey, K.K., Montoro, D.T., Mohan, R., Gazal, S., Engreitz, J.M., Xavier, R.J., Price, A.L., and Regev, A. (2022). Identifying disease-critical cell types and cellular processes by integrating single-cell RNA-sequencing and human genetics. Nat. Genet. 54, 1479–1492. 10.1038/s41588-022-01187-9.

11. Zhang, M.J., Hou, K., Dey, K.K., Sakaue, S., Jagadeesh, K.A., Weinand, K., Taychameekiatchai, A., Rao, P., Pisco, A.O., Zou, J., et al. (2022). Polygenic enrichment distinguishes disease associations of individual cells in single-cell RNA-seq data. Nat. Genet. 54, 1572–1580. 10.1038/s41588-022-01167-z.

12. de Leeuw, C.A., Mooij, J.M., Heskes, T., and Posthuma, D. (2015). MAGMA: generalized gene-set analysis of GWAS data. PLoS Comput. Biol. 11, e1004219. 10.1371/journal.pcbi.1004219.

13. Gupta, A., Weinand, K., Nathan, A., Sakaue, S., Zhang, M.J., Accelerating Medicines Partnership RA/SLE Program and Network, Donlin, L., Wei, K., Price, A.L., Amariuta, T., et al. (2023). Dynamic regulatory elements in single-cell multimodal data implicate key immune cell states enriched for autoimmune disease heritability. Nat. Genet. 55, 2200–2210. 10.1038/s41588-023-01577-7.

14. Björck, □ke, and Golub, G.H. (1973). Numerical methods for computing angles between linear subspaces. Math. Comput. 27, 579–594. 10.1090/S0025-5718-1973-0348991-3.

15. Rhee, J.-K., Joung, J.-G., Chang, J.-H., Fei, Z., and Zhang, B.-T. (2009). Identification of cell cycle-related regulatory motifs using a kernel canonical correlation analysis. BMC Genomics 10 *Suppl 3*, S29. 10.1186/1471-2164-10-S3-S29.

16. Yan, J., Qiu, Y., Ribeiro Dos Santos, A.M., Yin, Y., Li, Y.E., Vinckier, N., Nariai, N., Benaglio, P., Raman, A., Li, X., et al. (2021). Systematic analysis of binding of transcription factors to noncoding variants. Nature 591, 147–151. 10.1038/s41586-021-03211-0.

17. Heidersbach, A.J., Dorighi, K.M., Gomez, J.A., Jacobi, A.M., and Haley, B. (2023). A versatile, high-efficiency platform for CRISPR-based gene activation. Nat. Commun. 14, 902. 10.1038/s41467-023-36452-w.

18. Li, J., Bushel, P.R., Chu, T., and Wolfinger, R.D. (2009). Principal Variance Components Analysis: Estimating Batch Effects in Microarray Gene Expression Data. In Wiley Series in Probability and Statistics, A. Scherer, ed. (Wiley), pp. 141–154. 10.1002/9780470685983.ch12.

19. Subramanian, A., Tamayo, P., Mootha, V.K., Mukherjee, S., Ebert, B.L., Gillette, M.A., Paulovich, A., Pomeroy, S.L., Golub, T.R., Lander, E.S., et al. (2005). Gene set enrichment analysis: a knowledge-based approach for interpreting genome-wide expression profiles. Proc. Natl. Acad. Sci. U. S. A. 102, 15545–15550. 10.1073/pnas.0506580102.

20. Liberzon, A., Birger, C., Thorvaldsdóttir, H., Ghandi, M., Mesirov, J.P., and Tamayo, P. (2015). The Molecular Signatures Database (MSigDB) hallmark gene set collection. Cell Syst. 1, 417–425. 10.1016/j.cels.2015.12.004.

21. GTEx Consortium, Laboratory, Data Analysis &Coordinating Center (LDACC)—Analysis Working Group, Statistical Methods groups—Analysis Working Group, Enhancing GTEx (eGTEx) groups, NIH Common Fund, NIH/NCI, NIH/NHGRI, NIH/NIMH, NIH/NIDA, Biospecimen Collection Source Site—NDRI, et al. (2017). Genetic effects on gene expression across human tissues. Nature 550, 204–213. 10.1038/nature24277.

22. Psarras, A., Wittmann, M., and Vital, E.M. (2022). Emerging concepts of type I interferons in SLE pathogenesis and therapy. Nat. Rev. Rheumatol. 18, 575–590. 10.1038/s41584-022-00826-z.

23. GTEx Consortium (2020). The GTEx Consortium atlas of genetic regulatory effects across human tissues. Science 369, 1318–1330. 10.1126/science.aaz1776.

24. Heng, T.S.P., Painter, M.W., and Immunological Genome Project Consortium (2008). The Immunological Genome Project: networks of gene expression in immune cells. Nat. Immunol. 9, 1091–1094. 10.1038/ni1008-1091.

25. Painter, M.W., Davis, S., Hardy, R.R., Mathis, D., Benoist, C., and Immunological Genome Project Consortium (2011). Transcriptomes of the B and T lineages compared by multiplatform microarray profiling. J. Immunol. 186, 3047–3057. 10.4049/jimmunol.1002695.

26. Ota, M., Nagafuchi, Y., Hatano, H., Ishigaki, K., Terao, C., Takeshima, Y., Yanaoka, H., Kobayashi, S., Okubo, M., Shirai, H., et al. (2021). Dynamic landscape of immune cell-specific gene regulation in immune-mediated diseases. Cell 184, 3006–3021.e17. 10.1016/j.cell.2021.03.056.

27. Roadmap Epigenomics Consortium, Kundaje, A., Meuleman, W., Ernst, J., Bilenky, M., Yen, A., Heravi-Moussavi, A., Kheradpour, P., Zhang, Z., Wang, J., et al. (2015). Integrative analysis of 111 reference human epigenomes. Nature 518, 317–330. 10.1038/nature14248.

28. Zhang, F., Jonsson, A.H., Nathan, A., Millard, N., Curtis, M., Xiao, Q., Gutierrez-Arcelus, M., Apruzzese, W., Watts, G.F.M., Weisenfeld, D., et al. (2023). Deconstruction of rheumatoid arthritis synovium defines inflammatory subtypes. Nature 623, 616–624. 10.1038/s41586-023-06708-y.

29. Rubtsov, A.V., Rubtsova, K., Fischer, A., Meehan, R.T., Gillis, J.Z., Kappler, J.W., and Marrack, P. (2011). Toll-like receptor 7 (TLR7)-driven accumulation of a novel CD11c^+^ B-cell population is important for the development of autoimmunity. Blood 118, 1305–1315. 10.1182/blood-2011-01-331462.

30. Jonsson, A.H., Zhang, F., Dunlap, G., Gomez-Rivas, E., Watts, G.F.M., Faust, H.J., Rupani, K.V., Mears, J.R., Meednu, N., Wang, R., et al. (2022). Granzyme K+ CD8 T cells form a core population in inflamed human tissue. Sci. Transl. Med. 14, eabo0686. 10.1126/scitranslmed.abo0686.

31. Ziffra, R.S., Kim, C.N., Ross, J.M., Wilfert, A., Turner, T.N., Haeussler, M., Casella, A.M., Przytycki, P.F., Keough, K.C., Shin, D., et al. (2021). Single-cell epigenomics reveals mechanisms of human cortical development. Nature 598, 205–213. 10.1038/s41586-021-03209-8.

32. Nagy, C., Maitra, M., Tanti, A., Suderman, M., Théroux, J.-F., Davoli, M.A., Perlman, K., Yerko, V., Wang, Y.C., Tripathy, S.J., et al. (2020). Single-nucleus transcriptomics of the prefrontal cortex in major depressive disorder implicates oligodendrocyte precursor cells and excitatory neurons. Nat. Neurosci. 23, 771–781. 10.1038/s41593-020-0621-y.

33. Merkle, F.T., Tramontin, A.D., García-Verdugo, J.M., and Alvarez-Buylla, A. (2004). Radial glia give rise to adult neural stem cells in the subventricular zone. Proc. Natl. Acad. Sci. U. S. A. 101, 17528–17532. 10.1073/pnas.0407893101.

34. Ishigaki, K., Sakaue, S., Terao, C., Luo, Y., Sonehara, K., Yamaguchi, K., Amariuta, T., Too, C.L., Laufer, V.A., Scott, I.C., et al. (2022). Multi-ancestry genome-wide association analyses identify novel genetic mechanisms in rheumatoid arthritis. Nat. Genet. 54, 1640–1651. 10.1038/s41588-022-01213-w.

35. Mouri, K., Guo, M.H., de Boer, C.G., Lissner, M.M., Harten, I.A., Newby, G.A., DeBerg, H.A., Platt, W.F., Gentili, M., Liu, D.R., et al. (2022). Prioritization of autoimmune disease-associated genetic variants that perturb regulatory element activity in T cells. Nat. Genet. 54, 603–612. 10.1038/s41588-022-01056-5.

36. Kono, M., Hatano, H., Asahara, K., Nakano, M., Bagherzadeh, R., Kawashima, T., Arakawa, T., Sato, M., Inokuchi, H., Nishino, T., et al. (2025). Accurate, sensitive, and efficient chromatin accessibility quantification at target loci using UNIChro-seq. Preprint, 10.1101/2025.07.29.25332340 https://doi.org/10.1101/2025.07.29.25332340.

37. Robertson, C.C., Inshaw, J.R.J., Onengut-Gumuscu, S., Chen, W.-M., Santa Cruz, D.F., Yang, H., Cutler, A.J., Crouch, D.J.M., Farber, E., Bridges, S.L., et al. (2021). Fine-mapping, trans-ancestral and genomic analyses identify causal variants, cells, genes and drug targets for type 1 diabetes. Nat. Genet. 53, 962–971. 10.1038/s41588-021-00880-5.

38. Onengut-Gumuscu, S., Chen, W.-M., Burren, O., Cooper, N.J., Quinlan, A.R., Mychaleckyj, J.C., Farber, E., Bonnie, J.K., Szpak, M., Schofield, E., et al. (2015). Fine mapping of type 1 diabetes susceptibility loci and evidence for colocalization of causal variants with lymphoid gene enhancers. Nat. Genet. 47, 381–386. 10.1038/ng.3245.

39. Xie, G., Chen, X., Gao, Y., Yang, M., Zhou, S., Lu, L., Wu, H., and Lu, Q. (2025). Age-Associated B Cells in Autoimmune Diseases: Pathogenesis and Clinical Implications. Clin. Rev. Allergy Immunol. 68, 18. 10.1007/s12016-025-09021-w.

40. Weyand, C.M., and Goronzy, J.J. (2003). Ectopic germinal center formation in rheumatoid synovitis. Ann. N. Y. Acad. Sci. 987, 140–149. 10.1111/j.1749-6632.2003.tb06042.x.

41. Kawakami, E., Nakaoka, S., Ohta, T., and Kitano, H. (2016). Weighted enrichment method for prediction of transcription regulators from transcriptome and global chromatin immunoprecipitation data. Nucleic Acids Res. 44, 5010–5021. 10.1093/nar/gkw355.

42. Langmead, B., and Salzberg, S.L. (2012). Fast gapped-read alignment with Bowtie 2. Nat. Methods 9, 357–359. 10.1038/nmeth.1923.

43. Zhang, Y., Liu, T., Meyer, C.A., Eeckhoute, J., Johnson, D.S., Bernstein, B.E., Nusbaum, C., Myers, R.M., Brown, M., Li, W., et al. (2008). Model-based analysis of ChIP-Seq (MACS). Genome Biol. 9, R137. 10.1186/gb-2008-9-9-r137.

44. Wang, C., Sun, D., Huang, X., Wan, C., Li, Z., Han, Y., Qin, Q., Fan, J., Qiu, X., Xie, Y., et al. (2020). Integrative analyses of single-cell transcriptome and regulome using MAESTRO. Genome Biol. 21, 198. 10.1186/s13059-020-02116-x.

45. Nakano, M., Ota, M., Takeshima, Y., Iwasaki, Y., Hatano, H., Nagafuchi, Y., Itamiya, T., Maeda, J., Yoshida, R., Yamada, S., et al. (2022). Distinct transcriptome architectures underlying lupus establishment and exacerbation. Cell 185, 3375–3389.e21. 10.1016/j.cell.2022.07.021.

46. Bourgon, R., Gentleman, R., and Huber, W. (2010). Independent filtering increases detection power for high-throughput experiments. Proc. Natl. Acad. Sci. U. S. A. 107, 9546–9551. 10.1073/pnas.0914005107.

47. Robinson, M.D., McCarthy, D.J., and Smyth, G.K. (2010). edgeR: a Bioconductor package for differential expression analysis of digital gene expression data. Bioinforma. Oxf. Engl. 26, 139–140. 10.1093/bioinformatics/btp616.

48. Ianevski, A., Giri, A.K., and Aittokallio, T. (2022). Fully-automated and ultra-fast cell-type identification using specific marker combinations from single-cell transcriptomic data. Nat. Commun. 13, 1246. 10.1038/s41467-022-28803-w.

49. Hao, Y., Hao, S., Andersen-Nissen, E., Mauck, W.M., Zheng, S., Butler, A., Lee, M.J., Wilk, A.J., Darby, C., Zager, M., et al. (2021). Integrated analysis of multimodal single-cell data. Cell 184, 3573–3587.e29. 10.1016/j.cell.2021.04.048.

50. Bielecki, P., Riesenfeld, S.J., Hütter, J.-C., Torlai Triglia, E., Kowalczyk, M.S., Ricardo-Gonzalez, R.R., Lian, M., Amezcua Vesely, M.C., Kroehling, L., Xu, H., et al. (2021). Skin-resident innate lymphoid cells converge on a pathogenic effector state. Nature 592, 128–132. 10.1038/s41586-021-03188-w.

51. Butler, A., Hoffman, P., Smibert, P., Papalexi, E., and Satija, R. (2018). Integrating single-cell transcriptomic data across different conditions, technologies, and species. Nat. Biotechnol. 36, 411–420. 10.1038/nbt.4096.

52. International HapMap 3 Consortium, Altshuler, D.M., Gibbs, R.A., Peltonen, L., Altshuler, D.M., Gibbs, R.A., Peltonen, L., Dermitzakis, E., Schaffner, S.F., Yu, F., et al. (2010). Integrating common and rare genetic variation in diverse human populations. Nature 467, 52–58. 10.1038/nature09298.

53. Bycroft, C., Freeman, C., Petkova, D., Band, G., Elliott, L.T., Sharp, K., Motyer, A., Vukcevic, D., Delaneau, O., O’Connell, J., et al. (2018). The UK Biobank resource with deep phenotyping and genomic data. Nature 562, 203–209. 10.1038/s41586-018-0579-z.

54. Gazal, S., Finucane, H.K., Furlotte, N.A., Loh, P.-R., Palamara, P.F., Liu, X., Schoech, A., Bulik-Sullivan, B., Neale, B.M., Gusev, A., et al. (2017). Linkage disequilibrium-dependent architecture of human complex traits shows action of negative selection. Nat. Genet. 49, 1421–1427. 10.1038/ng.3954.

55. Yu, G., Wang, L.-G., Han, Y., and He, Q.-Y. (2012). clusterProfiler: an R package for comparing biological themes among gene clusters. Omics J. Integr. Biol. 16, 284–287. 10.1089/omi.2011.0118.

56. Wang, G., Sarkar, A., Carbonetto, P., and Stephens, M. (2020). A simple new approach to variable selection in regression, with application to genetic fine mapping. J. R. Stat. Soc. Ser. B Stat. Methodol. 82, 1273–1300. 10.1111/rssb.12388.

57. Hayashi, T., Ozaki, H., Sasagawa, Y., Umeda, M., Danno, H., and Nikaido, I. (2018). Single-cell full-length total RNA sequencing uncovers dynamics of recursive splicing and enhancer RNAs. Nat. Commun. 9, 619. 10.1038/s41467-018-02866-0.

58. Pierson, E., GTEx Consortium, Koller, D., Battle, A., Mostafavi, S., Ardlie, K.G., Getz, G., Wright, F.A., Kellis, M., Volpi, S., et al. (2015). Sharing and Specificity of Co-expression Networks across 35 Human Tissues. PLoS Comput. Biol. 11, e1004220. 10.1371/journal.pcbi.1004220.

59. Clement, K., Rees, H., Canver, M.C., Gehrke, J.M., Farouni, R., Hsu, J.Y., Cole, M.A., Liu, D.R., Joung, J.K., Bauer, D.E., et al. (2019). CRISPResso2 provides accurate and rapid genome editing sequence analysis. Nat. Biotechnol. 37, 224–226. 10.1038/s41587-019-0032-3.

