## Supplementary Note and Figures for "CATaN maps gene regulatory programs that shape genetic risk across complex diseases"

### Supplemental Note

#### CATaN parameter settings

To determine the optimal weighting scheme for constructing the TF-GRN matrix, we compared a binary encoding (presence or absence of TF binding) against exponential distance-based weighting (equation (1)) across a range of half-decay distances (1 kb, 5 kb, 10 kb, 20 kb, 100 kb) (Figures S1B). For each condition, we applied CCA between the resulting TF-GRN matrix and the transcriptome matrix and quantified the strength of the extracted shared variation using the singular values (SVs) returned by CCA. Because all downstream analyses in this study were based on the first ten CC components, we summed the corresponding first ten SVs (SV1–SV10) for each condition and used this sum as the comparison metric. Across all tested conditions, exponential weighting with a 10 kb half-decay distance produced the highest summed SVs (Figure S1C), indicating that this scheme most effectively captured the shared variation between the TF-GRN matrix and the transcriptome matrix. A 10 kb half-decay distance also extends beyond proximal promoters and encompasses regulatory interactions consistent with enhancer-mediated regulation; notably, this corresponds to the half-decay distance recommended for enhancer-driven regulation and used as the default for scATAC-seq in MAESTRO. We therefore used exponential weighting with a 10 kb half-decay distance for all analyses.

#### Transcriptome and TF gene projections

We generated two types of gene-level projections: TF gene-level projections and transcriptome gene-level projections (see Methods). Because these two projections are correlated (Figure S1D), we present only the transcriptome gene-level projections to avoid redundancy.

**Key biological insights revealed by CATaN (other than GTEx)**

Across the eight datasets analyzed here—spanning bulk and single-cell transcriptomes, human and mouse, and diverse immune cell contexts—several consistent patterns emerged. First, CC annotations enriched for autoimmune disease heritability (RA, SLE, Multiple Sclerosis (MS), T1D, Primary Biliary Cholangitis (PBC) and Inflammatory Bowel Disease (IBD)) were consistently associated with a core set of immune-related TFs, including RELA, STAG1, EP300, and SPI1, and with B cell transcriptional programs driven by PAX5 and IRF4. Second, these patterns were observed in both human and mouse datasets, supporting the cross-species relevance of the identified regulatory programs. Third, in several datasets, distinct CC axes captured separable disease associations: for example, in the cytokine CD4 dataset, CC3 and CC4 showed differential enrichment for T1D versus IBD and PBC, suggesting that CATaN can distinguish disease-specific regulatory components within a shared immune landscape. Below, we describe the results for each dataset in detail.

**[human immune cell dataset] (Table S7)**

CC2-bottom captured the primary autoimmune heritability signal, associated with B cell regulators and B cell transcriptional programs. CC1-bottom captured a distinct myeloid program.

- **Heritability:** CC2-bottom was enriched for MS, RA, T1D, celiac disease, SLE, and PBC.
- **TF:** CC2-bottom is enriched for PAX5, IRF4, SPI1, RELA, and STAG1. CC1-bottom was enriched for

myeloid regulators CEBPB and FOS.

- **Tr**: CC2-bottom was enriched for B cells (Naïve B, unswitched memory B, and double-negative B cells). CC1-bottom was enriched for neutrophils and low-density granulocytes.
- **Gene**: CC2-bottom was enriched for B cell-specific pathways. CC1-bottom was enriched for tuberculosis response and lysosomal processes.

**[ImmGen] (Table S8)**

Three CC axes captured autoimmune heritability in the mouse immune cell dataset, each associated with distinct TF and cell-type programs.

- **Heritability**: CC2-bottom was enriched for MS, RA, PBC, SLE, celiac disease, IBD, and asthma. CC4-bottom was enriched for MS and RA. CC5-bottom was enriched for MS, RA, PBC and SLE.
- **TF**: CC2 bottom was enriched for SPI1, EP300 and RELA. CC4-bottom was enriched for ZNF274, RB, MYC, and BRD4. CC5-bottom was enriched for MYC and SPI1.
- **Tr**: CC2-bottom was enriched for B cells. CC4-bottom was enriched for B cells and T cells. CC5-bottom was enriched for macrophages, pro B cells and dendritic cells.
- **Gene**: CC4-bottom was enriched for ATP-dependent chromatin remodeling.CC5-bottom was enriched for osteoclast differentiation.

**[ILC dataset] (Table S9)**

Two CC axes were enriched for autoimmune heritability in mouse skin-resident innate lymphoid cells, one linked to immune regulation and one to cell proliferation.

- **Heritability**: CC4-bottom and CC6-bottom were enriched for autoimmune diseases.
- **TF**: CC4-bottom was enriched for STAG1, RELA, SPI1, VDR, and EP300. CC6-bottom was enriched for NFYA, E2F1, E2F4, RB and MYB.
- **Tr**: CC4-bottom was enriched for ILC3. CC6-bottom was enriched for Birc5-high ILC.
- **Gene**: CC4-bottom was enriched for IBD and Th17 cell differentiation. CC6-bottom was enriched for DNA replication and cell cycle.

**[cytokine CD4 dataset] (Table S10)**

Three CC axes captured autoimmune heritability under cytokine stimulation conditions, with CC3-bottom and CC4-bottom resolving disease-specific regulatory components.

- **Heritability:** CC6-bottom was enriched for MS, RA, SLE, IBD, PBC, and T1D. CC3-bottom and CC4-bottom were both enriched for MS, RA, and SLE, but diverged in other traits: CC3-bottom was enriched for T1D, whereas CC4-bottom was enriched for IBD and PBC.
- **TF**: CC3-bottom was enriched for RELA and EP300. CC4-bottom was enriched for RELA and STAG1.
- **Tr**: CC3-bottom and CC4-bottom were both enriched for Th1-like stimulation conditions (CD3/28 stimulation, IL-12, anti-IL4).
- **Gene**: CC3-bottom was enriched for viral infection pathway. CC4-bottom was enriched for cytokine-related annotations.

**[Multi-autoimmune dataset] (Table S11)**

Cell-type-specific CATaN analysis identified distinct CC axes for RA and SLE, both converging on interferon-related pathways but differing in Tr enrichment patterns.

- **Heritability:** CC4-top of double-negative B cell (DNB) showed highest enrichment for RA. CC3-bottom of Th17 cell showed highest enrichment for SLE.
- **TF**: CC4-top (DNB) was enriched for RELA, STAG1, STAT1, EP300 and VDR. CC3-bottom (Th17 cells) was enriched for RELA, STAT1 and VDR.
- **Tr**: CC4-top (DNB) was enriched for SLE samples. CC3-bottom (Th17 cells) was not enriched for any specific disease.
- **Gene**: Both CC4-top (DNB) and CC3-bottom (Th17 cells) were enriched for interferon pathways.

**[RA synovium dataset]**

**<u>All cell-types dataset</u> (Table S12)**

CC1-top showed the broadest autoimmune heritability enrichment, with CC2-top capturing an overlapping but B cell-focused signal.

- **Heritability**: CC1-top was enriched for SLE, RA, IBD, MS, T1D, celiac disease, asthma, and PBC. CC2-top was enriched for SLE, RA, IBD, MS and T1D.
- **TF**: CC1-top was enriched for MYB and SPI1. CC2-top was enriched for SPI1, STAG1, RELA and EP300.
- **Tr**: CC1-top was enriched for B cells and T cells. CC2-top was enriched for B cells.
- **Gene**: CC1-top was enriched for interferon (IFN)-$\gamma$ response pathway and tumor necrosis factor-α (TNF-$\alpha$) signaling pathway. CC2-top was enriched for hematopoietic B cell receptor signaling pathway and cell lineage pathway.

**T cell dataset (Table S13)**

Four axes were enriched for RA heritability. These are enriched for GZMK/B$^+$ memory CD8$^+$ T cells and T peripheral helper (Tph) cells.

- **Heritability**: CC3-bottom, CC4-bottom, CC5-bottom and CC8-top were enriched for RA heritability.
- **TF**: CC3-bottom was enriched for MYC, GABPA, RB, E2F4, E2F1, and SIX5. CC4-bottom was enriched for AR and FOXA1. CC5-bottom was enriched for MAZ. CC8-top was enriched for E2F7, RBL2, RB, E2F4, E2F1, RELA and EP300.
- **Tr:** CC3-bottom was enriched for Tph. CC4-bottom was enriched for proliferating T cells, CD38$^+$ T cells and Tph. CC5-bottom was enriched for proliferating T cells and GZMK/B$^+$ memory CD8$^+$ T cells. CC8-top was enriched for proliferating T cells.
- **Gene:** CC3-bottom was enriched for TNFa signaling. CC4-bottom was enriched for Cell adhesion molecule interaction signaling and IFN-$\gamma$ response pathway. CC5-bottom was enriched for IFNα and γ response pathways and antigen processing and presentation pathway. CC8-top was enriched for E2F targets pathway, G2M checkpoint pathway and cell cycle pathway.

**B cell dataset (Table S14)**

Four axes were enriched for RA heritability. These were enriched for B cell populations such as autoimmune-associated B cells (ABCs) and germinal center-like (GC-like) B cells, the latter suggesting a role for ectopic germinal center formation in RA.

- **Heritability**: CC1-bottom, CC2-top and CC6-top and CC9-bottom were enriched for RA heritability.

- **TF**: CC1-bottom was enriched for CHD1, SPI1, ETS1, and RELA. CC2-top was enriched for SPI1 and ZNF274. CC6-top was enriched for MYB, STAG1, and CTCF. CC9-bottom was enriched for CHD1, AFF4, VDR, MED1 and MYC.
- **Tr:** CC1-bottom was enriched for GC-like B cells. CC2-top was enriched for ABCs. CC6-top and CC9-bottom were enriched for HLA-DR+IgG+ plasmablast and GC-like B cells.
- **Gene:** CC1-bottom was enriched for ribosome and autoimmune disease related pathways (including autoimmune thyroid disease and type1D). CC2-top was enriched for B cell receptor signaling. CC6-top was enriched for IFNγ response pathway. CC9-bottom was enriched for ribosome pathway.

**[Neural organoid dataset] (Table S15)**

Five CC axes were enriched for neurological disease heritability in neural organoid cells. Each of these axes highlighted distinct TF-GRNs associated with different biological functions, such as synaptic vesicle cycle and mismatch repair.

- **Heritability**: CC1-bottom, CC2-top, CC3-bottom, CC6-top and CC7-top were enriched for neurological diseases**.**
- **TF**: CC1-bottom was enriched for IRF5, ZNF274, SMAD2, AR (androgen receptor) and REST. CC2-top was enriched for CTCF. CC3-bottom was enriched for REST and GABPA. CC6-top was enriched for E2F4, E2F1 and KDM5B. CC7-top was enriched for REST, JARID2, and EZH2.
- **Tr**: CC2-top was enriched for radial glial cells and mature neurons. CC3-bottom was enriched for mature neurons. CC6-top was enriched for radial glial cells and neural stem cells. CC7-top was enriched for mature

neurons.

- **Gene**: CC1-bottom was enriched for axon guidance. CC2-top was enriched for reactive oxygen species and pathways of neurodegeneration. CC3-bottom was enriched for amyotrophic lateral sclerosis (ALS) pathway and synaptic vesicle cycle. CC6-top was enriched for DNA replication and mismatch repair. CC7-top was enriched for motor proteins.
-

#### Supplementary Figures

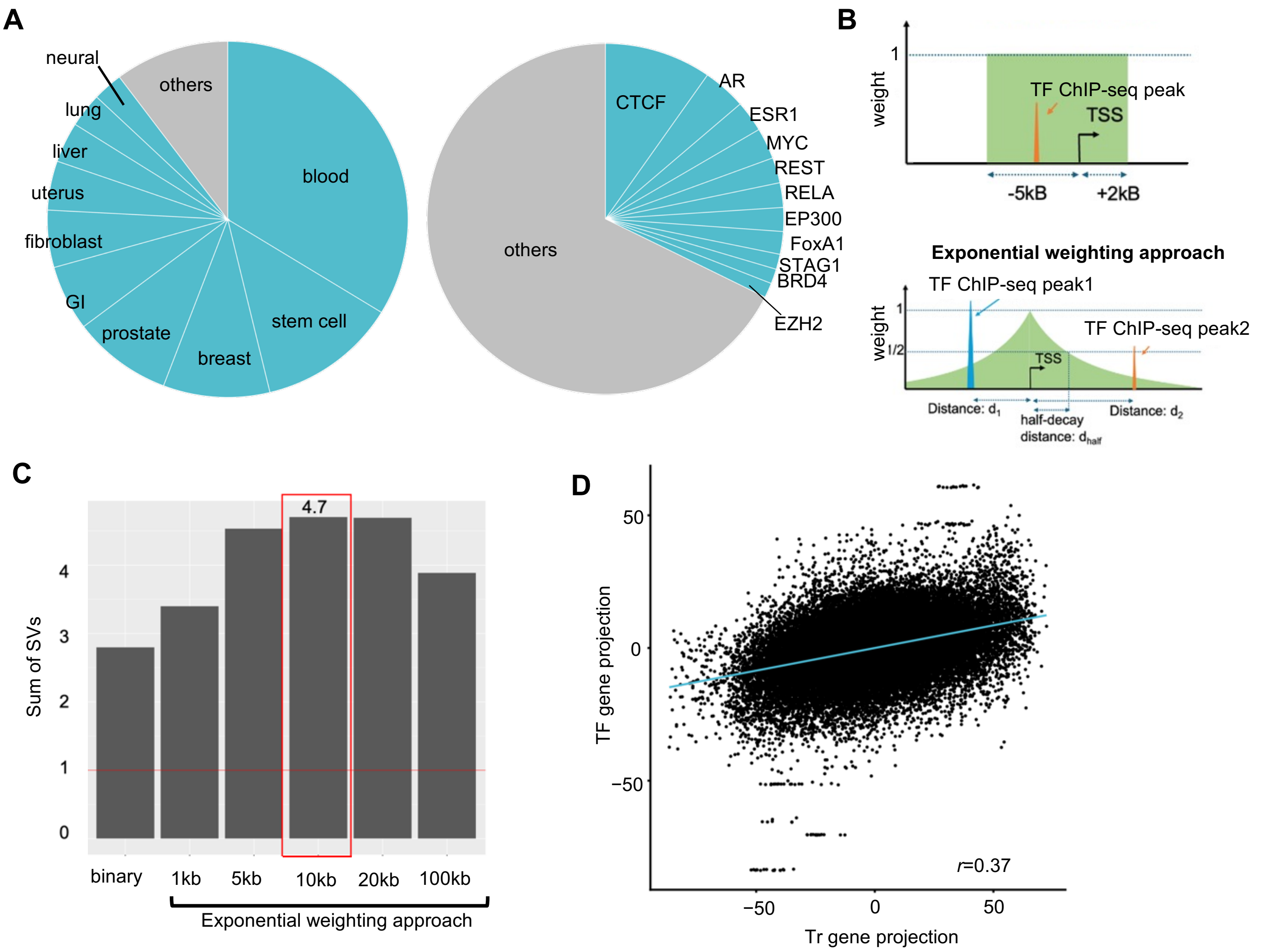


**Fig. S1 Composition of the TF-GRN matrix and parameter setting for CATaN.**

(A) Composition of the TF ChIP-seq dataset used to construct the TF-GRN matrix. Left, breakdown by tissue of origin; right, breakdown by target TF. GI, gastrointestinal tract.

(B) Schematic of two approaches for constructing the TF-GRN matrix: binary approach (top) and exponential weighting approach (bottom). TSS, transcription start site.

(C) Sum of singular values (SVs) from CC1–CC10 adjusted by null dataset across different half-decay distances ($d_{half}$), computed using the GTEx dataset as the transcriptome matrix.

(D) Comparison of transcriptome (Tr) gene projections (x axis) and TF gene projections (y axis) (n = 5,440) computed using the GTEx dataset as the transcriptome matrix. CC1 is shown as a representative result. r, Pearson correlation coefficient. The blue line indicates linear regression fit.

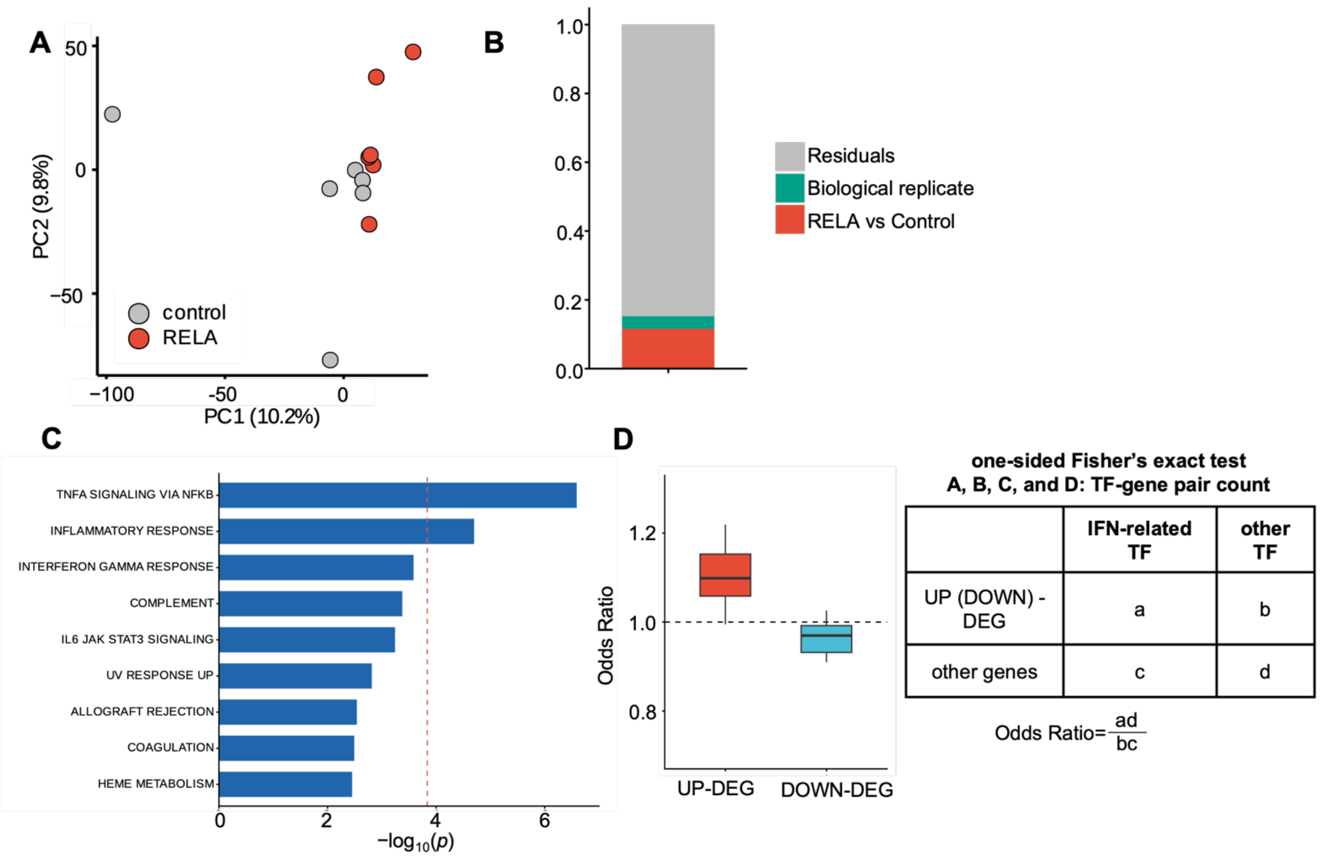


**Fig. S2 Supporting analyses for directional consistency.**

(A) Principal component analysis (PCA) of CRISPRa RELA overexpression RNA-seq data. RELA-overexpressed samples are shown in red. Variance explained is shown in parentheses. Only day 7 data used for CATaN ($n$ = 6 for sgRELA and sgControl) were included.

(B) Principal variance component analysis (PVCA) of CRISPRa RELA overexpression RNA-seq data.

(C) Gene set enrichment analysis (GSEA) of CRISPRa-upregulated genes using MSigDB Hallmark gene sets. Horizontal bar plot showing the pathways ranked by $-\log_{10}(p)$. Only pathways with a nominal $p < 0.05$ are shown. The dashed red line indicates the Bonferroni-corrected significance threshold.

(D) Enrichment of SLE-DEGs near interferon (IFN)-related TF binding sites ($n$ = 27 cell types). For each TF–gene pair, genes were classified as SLE upregulated DEGs or not, and TFs were classified as IFN-related (STAT and IRF family) or not. Odds ratios from Fisher's exact test indicate whether upregulated (or downregulated) DEGs are preferentially associated with IFN-related TF binding sites compared to other TF binding sites. An odds ratio >1 indicates that DEGs are disproportionately found near IFN-related TFs rather than other TFs.

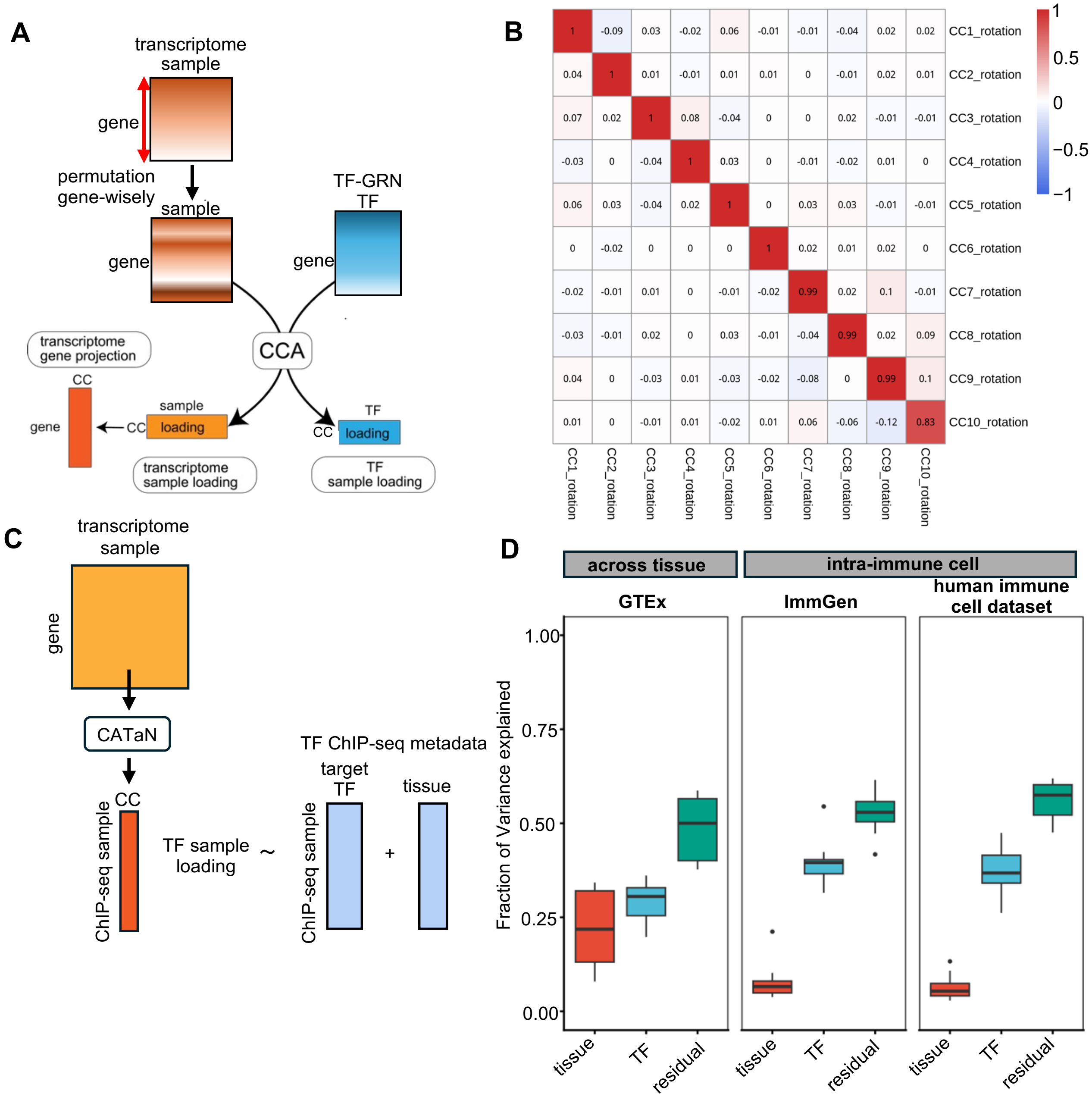


**Fig. S3 Robustness and sensitivity analysis of CATaN.**

(A) Schematic of the CATaN workflow and the permutation procedure used to construct the null model. To construct the null model, gene labels in the transcriptome matrix are permuted.

(B) Pearson correlation coefficients of TF sample loadings between two independent half-subsets of GTEx.

(C) Overview of variance partitioning of CATaN sample loadings. TF sample loadings (CC1–CC10) were regressed on TF ChIP-seq metadata to compute explained variance. The same procedure was applied to transcriptome sample loadings and transcriptome metadata.

(D) Variance partitioning of CATaN TF sample loadings (*n* = 10 CC axes) across three datasets: GTEx (across-tissue), ImmGen (intra-immune cell), and human immune cell dataset (intra-immune cell). Each box plot summarizes the proportion of variance explained by tissue, TF, and residual across the ten canonical components.

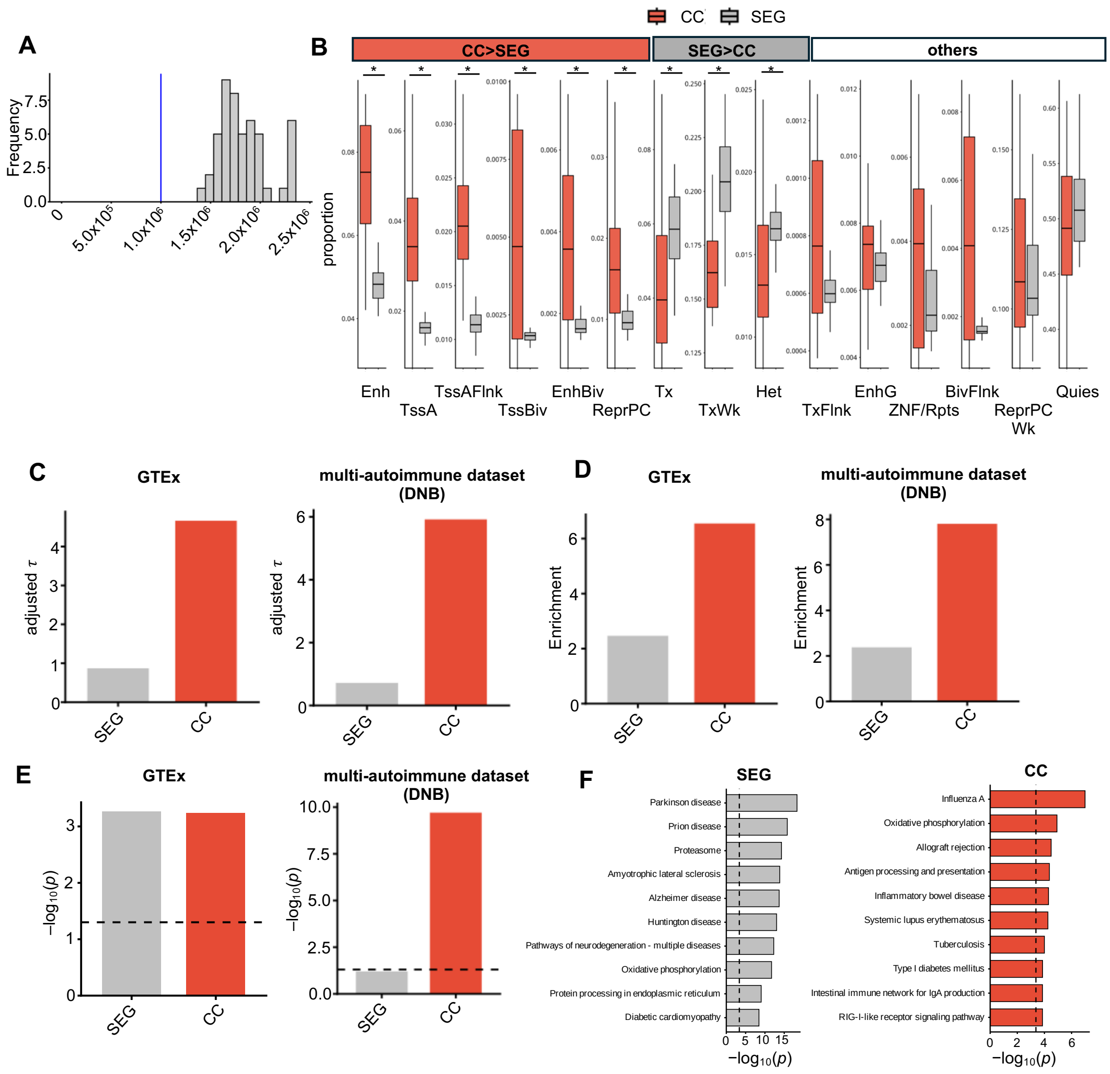


**Fig. S4 Comparison of CC and SEG annotations.**

(A) Number of SNPs included in each annotation. Because SEG annotations ($n$ = 53) varied in size across annotations, their distribution is shown as a histogram. The CC annotation size ($n$ = 20) was constant across annotations and is indicated by a blue vertical line. Transcriptome data: GTEx.

(B) Box plots showing the proportion of SNPs mapping to each chromatin state category defined by the Roadmap Epigenomics core 15-state ChromHMM model (SEG, n = 53; CC, n = 20). Chromatin state assignments were based on Roadmap sample E071 (brain). Asterisks denote categories with significant differences between CC and SEG annotations after Bonferroni correction (Wilcoxon rank-sum test). Chromatin states were defined using the 15-state ChromHMM model (Supplementary Table X).

(C–E) Comparison of CC and SEG annotations for a representative GWAS trait (RA, EUR). For each method, the annotation track with the highest $-\log_{10}(p)$ of $\tau$ is shown. Dashed lines indicate significance thresholds. (C) Adjusted $\tau$. (D) Enrichment. (E) Bonferroni-corrected $-\log_{10}(p)$ of $\tau$ for GTEx-based annotations (SEG, n = 54; CC, n = 20) and double-negative B cell (DNB) subset of the multi-autoimmune dataset (n = 20 each for SEG and CC). In **e,** dashed lines indicate nominal significance threshold ($-\log_{10}(0.05)$) applied to Bonferroni-corrected *p* values**.**

**(F)** Over-representation analysis (ORA) of DEG-based gene sets (top 1,000 upregulated DEGs by Z score) and CC transcriptome gene projections (top 10% by absolute CC score, *n* = 680) based on the double-negative B cell subset of the multi-autoimmune dataset. MSigDB Hallmark gene sets were used. The dashed line indicates the Bonferroni-corrected significance threshold. Enh, Enhancers; TssA, Active TSS; TssAFlnk, Flanking Active TSS; TssBiv, Bivalent/Poised TSS; EnhBiv, Bivalent Enhancer; ReprPC, Repressed PolyComb; Tx, Strong transcription; TxWk, Weak transcription; Het, Heterochromatin; TxFlnk, Transcr. at gene 5' and 3'; EnhG, Genic enhancers; ZNF/Rpts, ZNF genes & repeats; BivFlnk, Flanking Bivalent TSS/Enh; ReprPCWk, Weak Repressed PolyComb; Quies, Quiescent/Low.

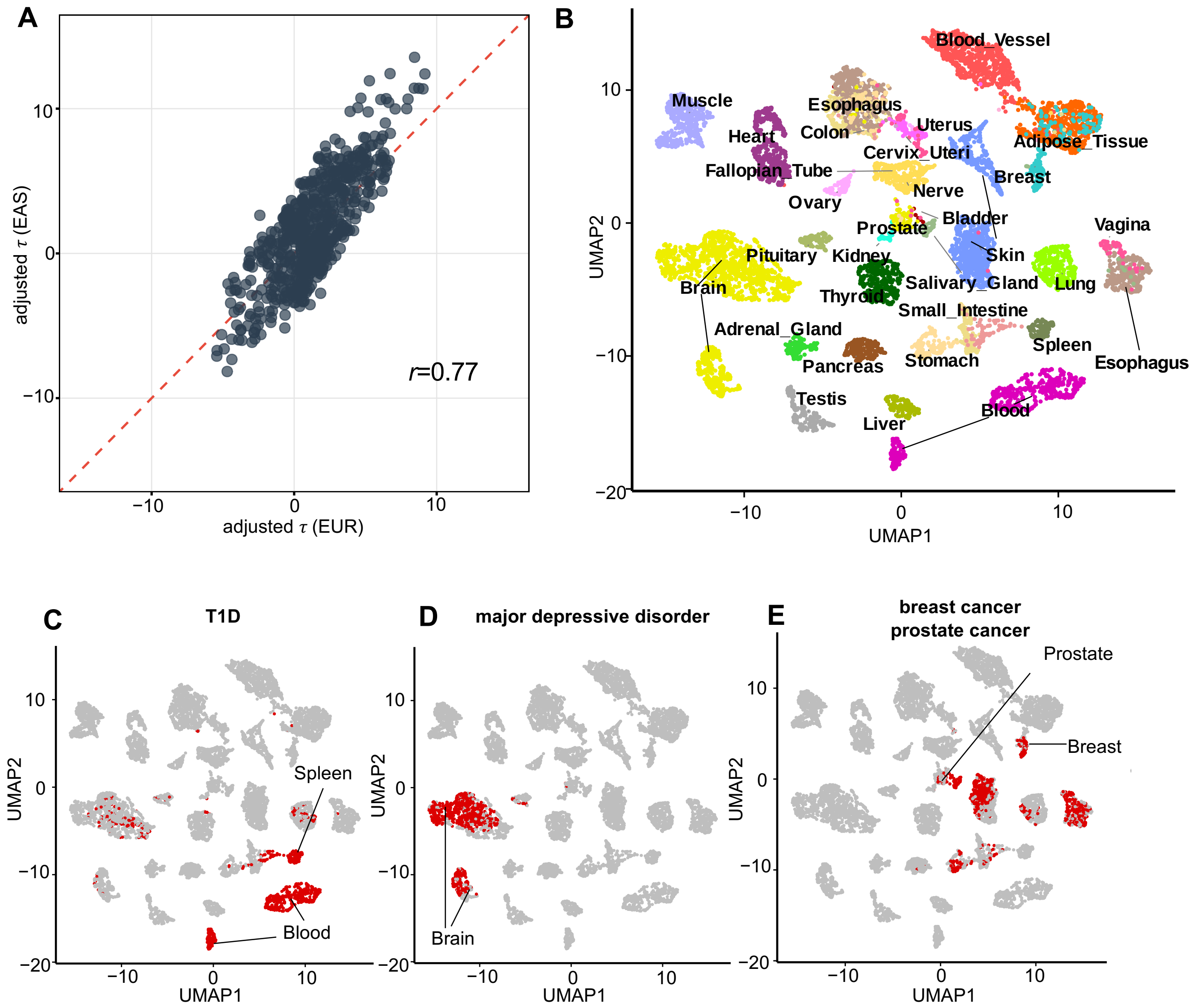


**Fig. S5 Cross-ancestry consistency and heritability enrichment mapped onto GTEx UMAP space.**

(A) Comparison of adjusted $\tau$ values between EUR (x axis) and EAS (y axis), GWAS trait: RA. The red dashed line indicates y = x.

(B–E) UMAP of GTEx dataset. (B) UMAP colored by GTEx tissue annotation. (C) Top 10% of Tr sample loadings from the CC annotation track most enriched for type 1 diabetes (T1D) heritability, highlighted in red. (D) As in C, for major depressive disorder (MDD). (E) As in C, for breast cancer and prostate cancer.

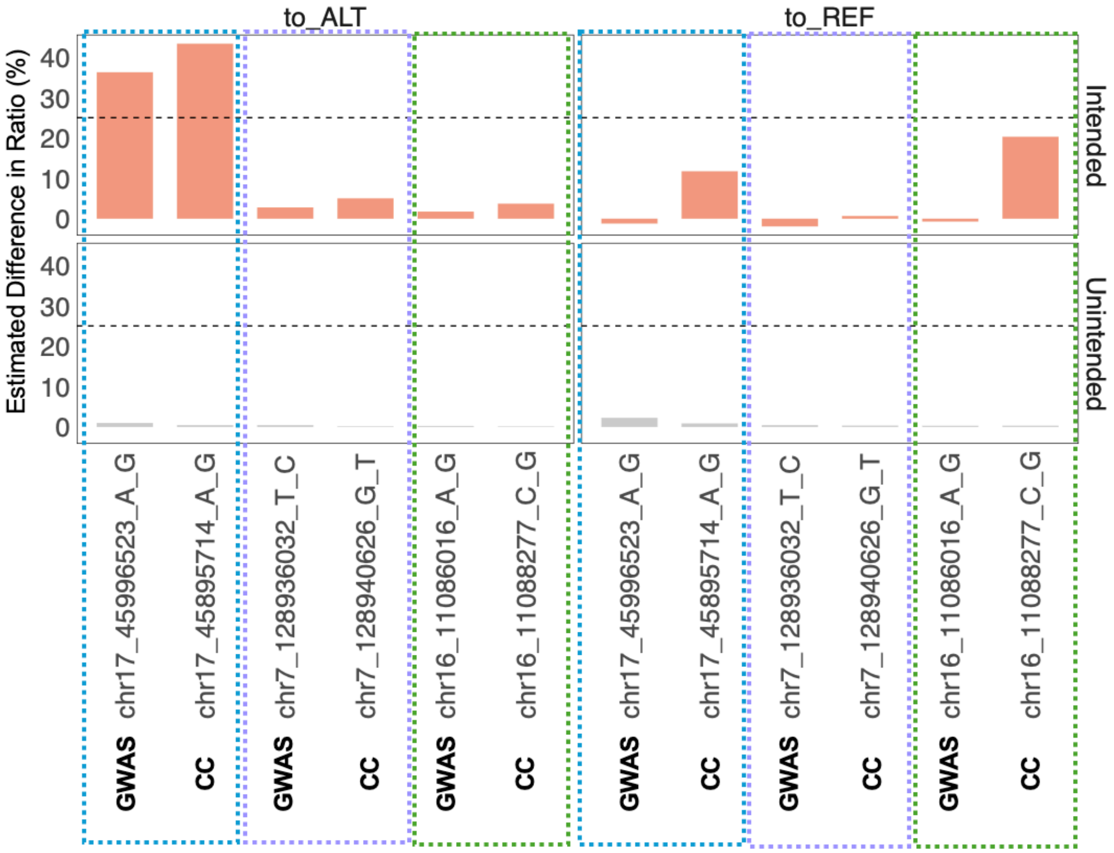


**Fig. S6 Genome editing efficiencies.**

Editing efficiencies of GWAS lead SNPs and their paired CC SNPs across three genomic regions (six SNPs in total). Boxes of the same color indicate SNP pairs (a GWAS lead SNP and its LD-partner CC SNP) from the same genomic region. Results are shown separately for editing toward the alternative (to_ALT) and reference (to_REF) alleles. Upper panels show intended editing efficiency (%); lower panels show unintended editing rate (%). The dashed line denotes the 25% inclusion threshold.
